# Optimization of universal primer set for the identification of all Influenza A viruses using real-time PCR

**DOI:** 10.1101/2022.08.30.505914

**Authors:** Altrice Harkness, Qiana L Matthews, Lula Smith, Bokai Roberston, Hongzhuan Wu

## Abstract

Avian Influenza Virus (AIV) occurs worldwide among aquatic wild and may mutate into a strain capable of efficient human-to-human transmission and cause a pandemic that could kill a large fraction of the human population. The purpose of this study is to efficiently identify and diagnose the virus for immediate treatment and prevention. Real-Time Reverse Transcription-Polymerase Chain Reaction (RRT-PCR or qPCR) was identified as the most efficient way to detect AIV in previous laboratory research, however only one pair of primers was used. In order to further optimize the test and make it commercially available in the future, we first tested an eight pair primer set via RRT-PCR by using standard AIV isolates from Dr.Gaimbrone’s lab (Auburn University). Using the AIV isolates, the optimal cDNA and primer concentrations were determined, this aided in the identification of the most sensitive primer set. Primers PA, HA, NP, and NA were able to detect the cDNA of positive AIV isolates at a low concentration of 0.0039 mg/ml. AIV cDNA were detected by primers M and HA at its lowest concentration of 0.0001 and 0.01 pmol/μl. In order to further test our lab results, 20 fecal samples from aquatic wild birds were collected from Alabama Shakespeare Park located in Montgomery, Alabama during the winter season. The samples were analyzed using RRT-PCR. HA primer set detected 10 positive samples for Avian Influenza and is the most sensitive primer set to be used in RRT-PCR. The field sample detection confirmed laboratory results and revealed that local aquatic birds are actively shedding low pathogenic avian influenza virus(LPAIV). The optimization of all primer sets used in this study will aid in disease surveillance, facilitate quick diagnosis of Influenza A viruses, and implement an effective prevention of flu pandemic.

## Introduction

Avian Influenza Virus Type A is an emerging public health concern due to its ability to produce major illnesses that could result in death among millions of humans and birds, if not properly detected and contained. It is important to implement effective testing mechanisms for early identification to avoid a major outbreak among poultry as well as human to human transmission. Due to mutations, AIV strains possess the capability to alternate between low pathogenicity (LP) and high pathogenicity (HP) through antigenic shift and drifts. This could result in the development of a deadly strain. The virus can potentially be transmitted to mammals generating increased chances to spread from human to human. Avian Influenza (AI) is shed naturally among wild aquatic birds and other bird species. However, the critical concern is the spread of AIV to domesticated poultry, eggs and other animal species, such as swine. Although AI cases among humans are rare it is indefinitely possible to contract and spread the virus (1). Between 1959 and 1995, health officials noticed an AIV emergence, there were only 15 cases among poultry and no human infections were reported (2). The virus has been on the rise since then causing major outbreaks in birds crediting wild migratory birds for the spread. Recent cases include viral transmission in Nepal, China, Taiwan and Vietnam. These cases have occurred through the ingestion of fresh duck blood, undercooked poultry, and other delicacies (3).

In the United States, the CDC recognized and confirmed over 200 cases of HPAIV among the domestic flock. This potential threat caused the CDC to release a health alert advisory which warned individuals who work with poultry to stay away from dead or infected fowl. Though rare, AI strains have caused human infection, virus subtypes H5, H7, and H9 are responsible for the majority of AIV cases. The varying subtypes could exchange genetic material and produce genetic assortments leading to a human-transmissible strain of AIV (4). Identifying the various antigenic and genetic differences of the Influenza Type A Virus could implement effective diagnosis and vaccines to obviate transmission. Consumption of undercooked, infected poultry, direct contact with infected birds’ secretions or objects could lead to a major outbreak triggering a pandemic. Human health and economical trade among poultry could be at risk if this occurs, therefore access to rapid diagnostic test will help monitor disease surveillance and facilitate treatment to infected individuals.

The increase in the occurrence of Avian Influenza worldwide in poultry and humans introduces a potential influenza A pandemic that could pose a threat to both human health and the global economy. The goal of my research is to help establish an efficient method to help identify and quickly diagnose Avian Influenza. To systematically identify and analyze the 18 HA and 9 NA subtypes of influenza A virus, we need reliable, simple methods that not only characterize partial sequences but analyze the entire influenza A genome. In the past ten years, our research group has been using real-time quantitative (RT-PCR) to identify LPAI circulating in wild aquatic birds in Southern US. The primer sets we used are designed for the HA genes of all AIV subtypes. We compared 8 universal primers designed for AIV in this study and aimed to find which one is the most sensitive primer using real-time quantitative RT-PCR. Optimization of primer used for RT-PCR will accelerate the diagnosis of AIV and create an accurate therapy method for treatment.

## MATERIALS AND METHODS

### A. Sampling

Sampling was conducted in late December to early February, from 2017 to 2018. Fecal samples were collected from nesting waterfowl from local parks in Alabama. Swabs were placed in a tube containing 1 µL of virus-transport medium Phosphate-Buffered Saline (PBS) and 10µL of 5% fetal bovine serum (FBS). Samples were kept cold in the field with wet ice or frozen in dry ice and transferred to a − 70°C freezer within 1h. Samples were removed from freezer and frozen tubes containing cloacal swabs in transport medium were thawed and centrifuged at 2,000 × g for 10 min.

### B. Extracting RNA from virus samples

Total RNA was isolated and extracted from fecal samples using TRIzol (Invitrogen, Carlsbad, CA) by following the RNA isolation protocol. 500 µL of each fecal samples were extracted from vials and added to separate vials with 500 µL of TRIzol. Phase separation was done by incubating the homogenized samples for 5 minutes at 15 to 30°C to complete dissociation of nucleoprotein complexes. 200 µL of chloroform was added to samples and caps were secured before shaking tube by hand for 15 seconds. The mixtures were then incubated at 15 to 30°C for 2 to 3 minutes. Samples were centrifuged at 12,000 × g for 10 minutes at 4°C. The mixture separated into a lower red, phenol-chloroform phase, and a colorless upper aqueous phase. The aqueous phase consists of RNA and this was extracted and transferred to a new tube. RNA was precipitated from aqueous phase by mixing with 500 µL of isopropyl alcohol. The supernatant was removed and the RNA pellet was washed with 200 µL of 70% ethanol. Samples were mixed by vortexing tube and was centrifuged at 7,500× g for 5 minutes at 4°C. The RNA was re-dissolved initially by air drying the pellet for 5 minutes. The RNA samples was dissolved in RNase-free water by passing the solution through a pipette and incubated for 10 minutes at 55°C. Samples were then stored in freezer at − 70°C. The RNA concentration of each sample was measured by using Spectrophotometer.

### C. Synthesis of cDNA

Protocol Superscript IV First-Strand cDNA Synthesis Reaction was followed in this procedure. The primer was annealed to template RNA. 1 µL of 50 ng/µL random hexamers, 1 µL of 10mM dNTP mix, 10 µL of template RNA samples, and 12 µL of DEPC-treated water were mixed and centrifuged. The RNA-primer mix was heated to 65°C for 5 minutes and incubated on ice for 1 minute.

The RT (real-time) reaction mix was prepared by adding 4 µL of 5× SSIV buffer, 1 µL of mM DTT, 1 µL RNaseOUT™ Recombinant RNase Inhibitor enzyme, 1µL SuperScript® IV Reverse Transcriptase (200 U/µL). The mixture was briefly centrifuged to mix. The annealed RNA and RT reaction mix was combined and the reactions were incubated for 10 minutes at room temperature (23°C). The reaction mixtures were then placed in a thermal cycler for 20 minutes that followed a program. While in the cycler the samples were incubated at 50-55°C for 10 minutes and the reaction was inactivated by incubating at 80°C for an additional 10 minutes. The samples were then stored at −20°C awaiting PCR amplification. cDNA samples concentration was measured using spectrophotometry.

### D. Real-Time Reverse Transcription PCR (RRT-PCR)

#### 1. Preparation of RRT-PCR with serial dilution of AIV

A serial dilution was prepared by assembling PCR amplification mix following the SuperScript® IV RT-PCR control reactions protocol. Positive Control AIV virus H10N7 (P3) strain was provided by Dr.Giambrone’s lab in Auburn University. The positive cDNA sample (P3) was diluted from 10^1^ −10^8^. Each positive cDna dilution was added to a PCR plate wells 1-8 and A-H with the following 10 µL of 2X guarantee SYBR Green PCR master mix, 1 µL of cDNA diluted sample 1 µL of Primers PB2-NS (Table 1), and 7 µL of Rnase-Free water. Each well contained approximately 20 µL. After the PCR plate was prepared results were analyzed using RRT-PCR. A negative control of 20 µL of water was added in well A9 and A10

**Table 1.**
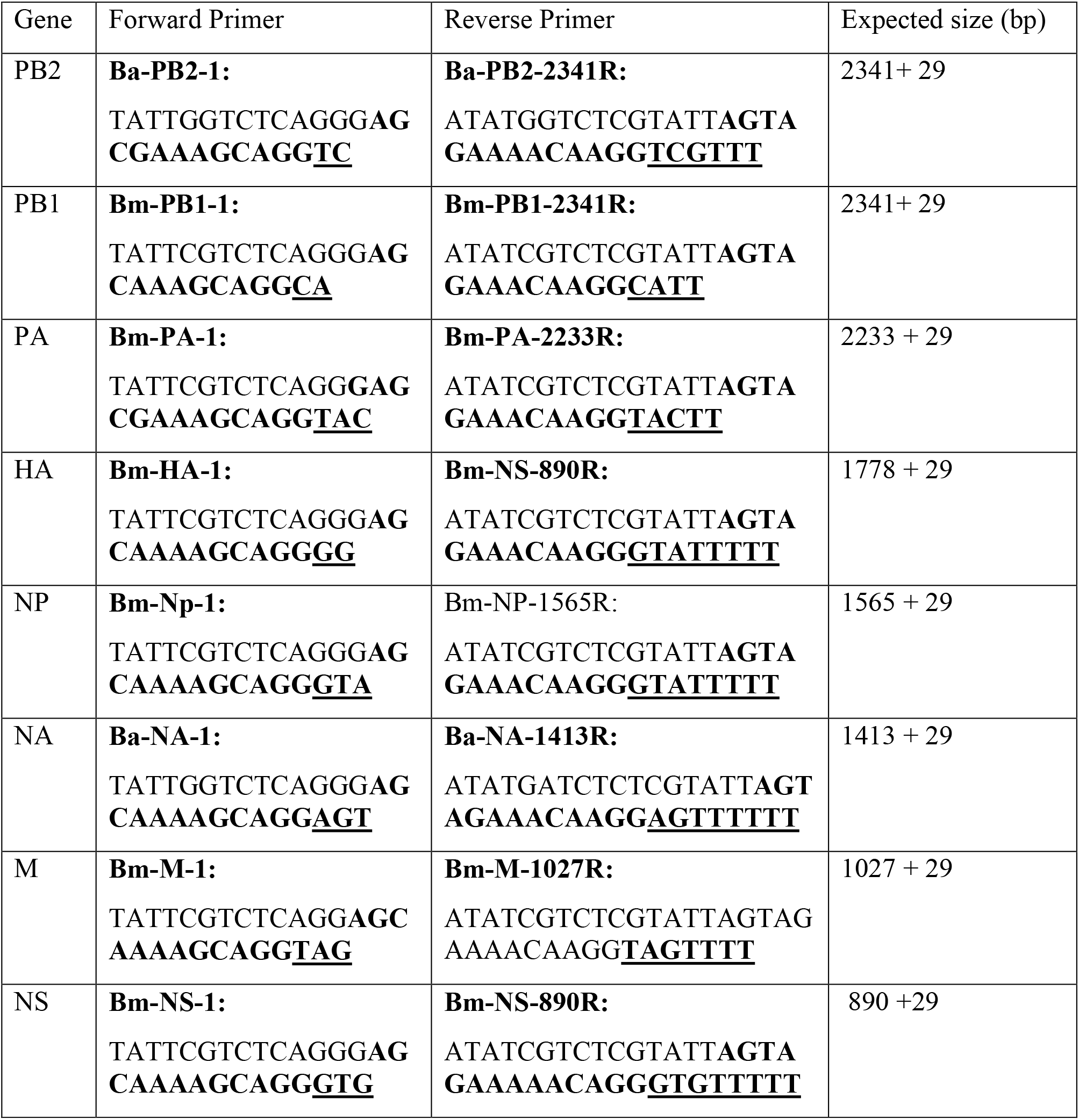
Primer set used for RT-PCR amplification of the eight vRNAs of Influenza A viruses.

#### 2. Preparation of RRT-PCR with serial dilution of universal primers(7)

A serial dilution was prepared by assembling PCR amplification mix following the SuperScript® IV RT-PCR control reactions protocol. Positive sample (P3) was diluted to 1:100. Each pair of primers were diluted from 10^1^ −10^8^ and the same protocol was followed. Negative Control was added in well A9 and A10.

#### 3. Optimization of RRT-PCR with 20 field samples and 8 primer Set(8)

PCR plate was prepared by adding AIV field samples with primer set and the SuperScript® IV RT-PCR control reactions protocol. Negative Control added in well A11 (20 µL of water) and positive control (P3 sample) in A12.

#### E. Experimental Design Outline

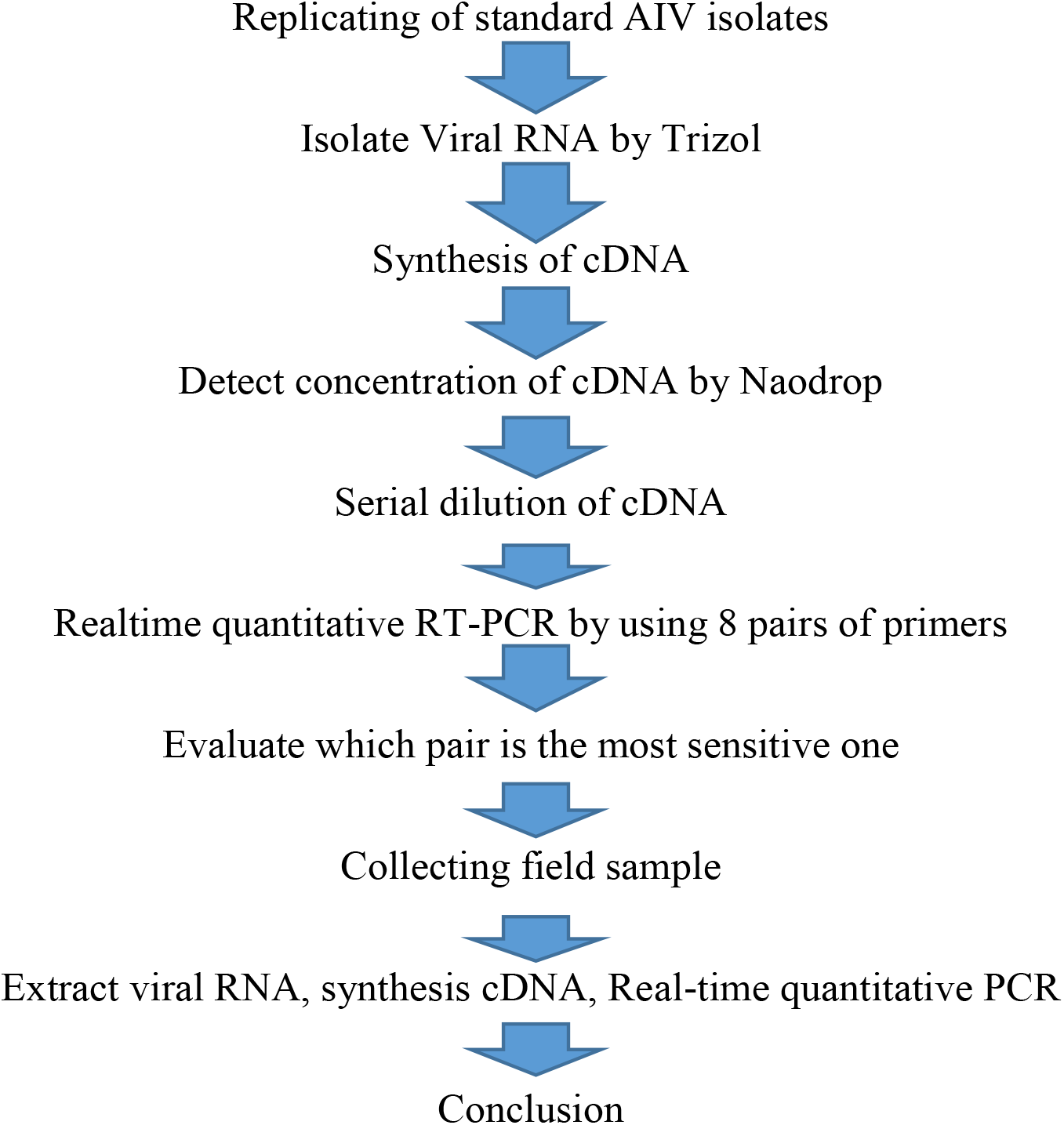

## RESULTS

### RNA Extraction/ cDNA synthesis

Forward and Reverse primer sets used in this research are shown in Table 1. The concentration of AIV viral RNA and cDNA field samples was detected via spectrophotometry shown in table 2 and table 3. The concentration of RNA collected from fecal samples gave a different absorbance at wavelengths of A=260. The highest concentration was field sample 19 at 39.49 mg/ml of RNA with an absorption of 0.987. Field sample 6 had the next highest concentration at 39.04 mg/ml with an optical density of 0.976. Field sample 7 had the lowest absorbance at 0.846 and corresponding RNA concentration at 33.85 mg/ml. Both positive controls, P3 and P4, each read over an absorbance of 1 with a concentration of 41 mg/ml. Each field sample served as a template in the next phase of the experiment for the synthesis of complementary DNA (cDNA).

**Table 2:** RNA Concentration of 20 field samples

| **RNA Samples** | **Concentration (mg/ml)** | **A260:** |
| --- | --- | --- |
| Field Sample 1 | 36.4109 | 0.91 |
| Field Sample 2 | 35.9582 | 0.9 |
| Field Sample 3 | 37.4217 | 0.936 |
| Field Sample 4 | 38.8843 | 0.972 |
| Field Sample 5 | 37.4442 | 0.936 |
| Field Sample 6 | 39.0425 | 0.976 |
| Field Sample 7 | 33.8535 | 0.846 |
| Field Sample 8 | 38.2446 | 0.956 |
| Field Sample 9 | 37.52680 | 0.938 |
| Field Sample 10 | 36.7894 | 0.92 |
| Field Sample 11 | 35.7524 | 0.894 |
| Field Sample 12 | 34.2108 | 0.855 |
| Field Sample 13 | 36.7822 | 0.92 |
| Field Sample 14 | 36.4816 | 0.912 |
| Field Sample 15 | 35.6777 | 0.892 |
| Field Sample 16 | 37.1394 | 0.928 |
| Field Sample 17 | 35.2034 | 0.88 |
| Field Sample 18 | 36.0059 | 0.9 |
| Field Sample 19 | 39.4865 | 0.987 |
| Field Sample 20 | 36.7317 | 0.918 |
| **RNA Positive Controls** |  |  |
| P3 | 41.7469 | 1.044 |
| P4 | 45.9558 | 1.149 |

Viral RNA concentrations and corresponding absorbances of field samples used in RRT-PCR.

**Table 3:** cDNA Concentration of 20 field samples

| **cDNA Samples** | **Concentration(mg/ml)** | **A260:** |
| --- | --- | --- |
| Field Sample 1 | 37.9544 | 1.026 |
| Field Sample 2 | 32.3391 | 0.874 |
| Field Sample 3 | 35.44904 | 0.959 |
| Field Sample 4 | 35.1718 | 0.951 |
| Field Sample 5 | 34.8523 | 0.942 |
| Field Sample 6 | 34.8172 | 0.941 |
| Field Sample 7 | 39.9169 | 1.079 |
| Field Sample 8 | 33.4272 | 0.903 |
| Field Sample 9 | 33.7455 | 0.912 |
| Field Sample 10 | 36.8773 | 0.997 |
| Field Sample 11 | 33.8773 | 0.916 |
| Field Sample 12 | 34.2386 | 0.925 |
| Field Sample 13 | 33.7784 | 0.913 |
| Field Sample 14 | 34.7892 | 0.94 |
| Field Sample 15 | 35.4174 | 0.957 |
| Field Sample 16 | 34.4634 | 0.931 |
| Field Sample 17 | 34.3132 | 0.927 |
| Field Sample 18 | 38.404 | 1.038 |
| Field Sample 19 | 34.32 | 0.928 |
| Field Sample 20 | 33.6539 | 0.91 |
| **cDNA Positive Controls** |  |  |
| P3 | 39.07689 | 1.075 |
| P4 | 39.511 | 1.068 |

cDNA concentrations and corresponding absorbance of field samples used in RRT-PCR.

Real-Time Reverse Transcription-PCR was used in synthesizing our cDNA for the extracted RNA in each field sample. The results of the synthesis are displayed in table 3. The yields of cDNA from the extracted RNA generally showed a lower absorbance. Idealistically cDNA samples concentrations will be lower when compared to RNA concentration. However, Field samples 1, 7 and 18 varied from normality having increased concentrations shown as in table 3. Each sample had a concentration of 37.96 mg/ml, 39.92 mg/ml and 38.404 mg/ml respectively, contrasting from lower RNA template concentrations of 36.41 mg/ml, 33.85 mg/ml and 36.01 mg/ml. The abnormalities could partially be due to the difference in concentrations of template RNA in each individual triplicate.

### RRT-PCR: The sensitivity of RRT-PCR with serial diluted AIV

The sensitivity of RRT-PCR was tested with AIV isolates from Dr.Giambrone’s laboratory (Figure 1). The detection thresholds of standard AIV with 8 pairs of universal primers could be detected by RRT-PCR listed in Figures 2-9. Briefly, cDNA was made ten-fold serial dilution from 1:10 to 1:10^8^ as template, real-time PCR was performed according to the description in the materials and methods. To measure qPCR data analysis, the relative expression levels of the specific gene in standard AIV samples were compared to that of negative samples and calculated using the formula 2^ΔΔCt^ where ΔΔCt =ΔCt (standard AIV)-ΔCt (negative control). Each analysis was performed in triplicate. Data were analyzed by analysis of variance (ANOVA) using Sigma Stat statistical analysis software (Systat Software, San Jose, CA). Statistical significance was set at P<0.05, results are summarized in table 4.

**Table 4:** Summarized detection sensitivity in RRT-PCR with 8 pairs of universal primer.

| Universal Primer | cDNA(mg/ml) |
| --- | --- |
| PB2 | 0.39 |
| PB1 | 0.39 |
| PA | 0.0039 |
| HA | 0.0039 |
| NP | 0.0039 |
| NA | 0.0039 |
| M | 3.9 |
| NS | 3.9 |

Optimal AIV cDNA concentrations measured with universal primer. Lowest concentration of 0.0039 mg/ml detected by primers PA, HA, NP, and NA.

**Figure 1.**
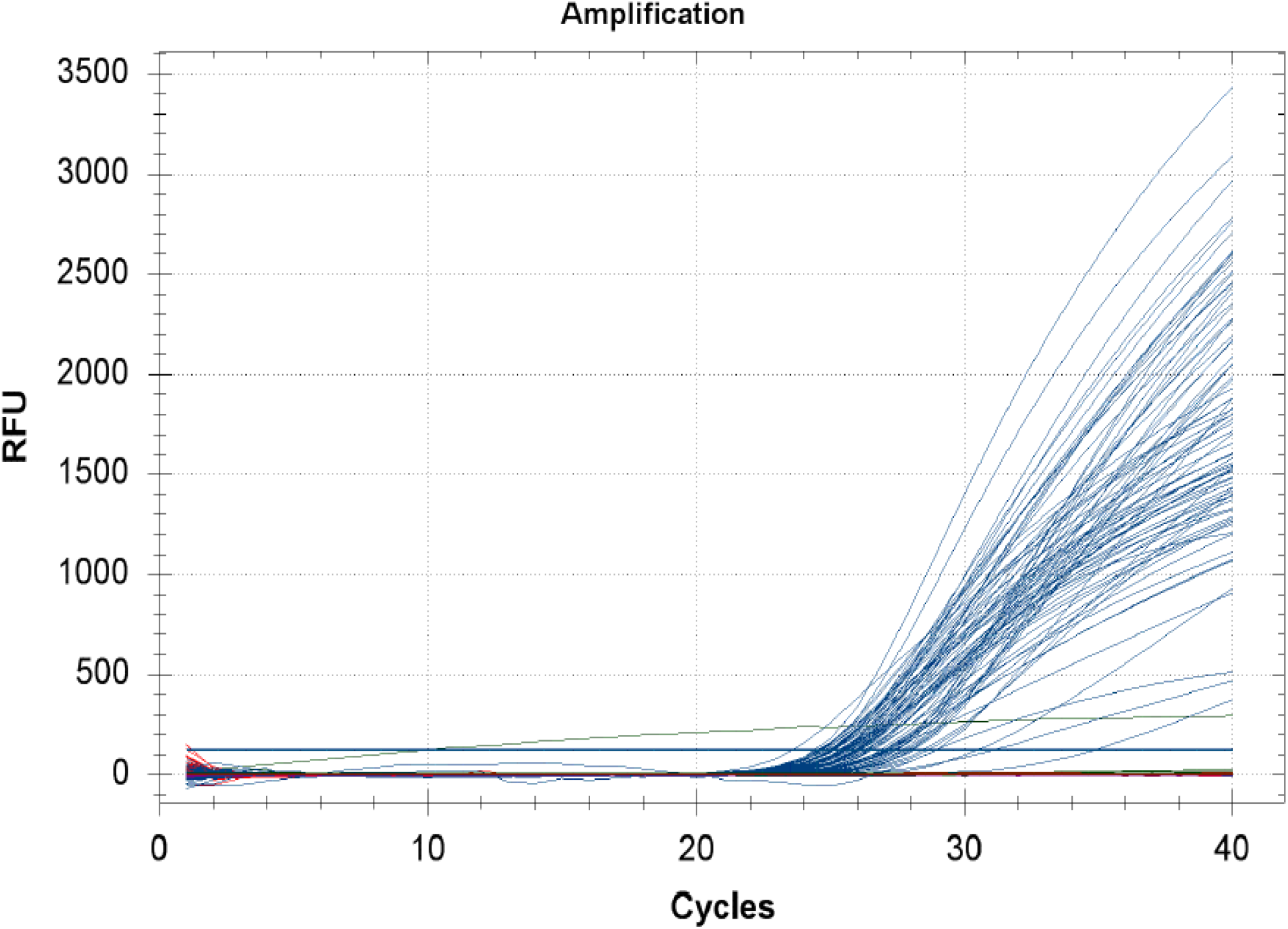
Sensitivity of RRT-PCR using serial diluted AIV. Relative Florence Units measured against cycles to determined amplification of positive diluted AIV Isolates with universal primers.

**Figure 2.**
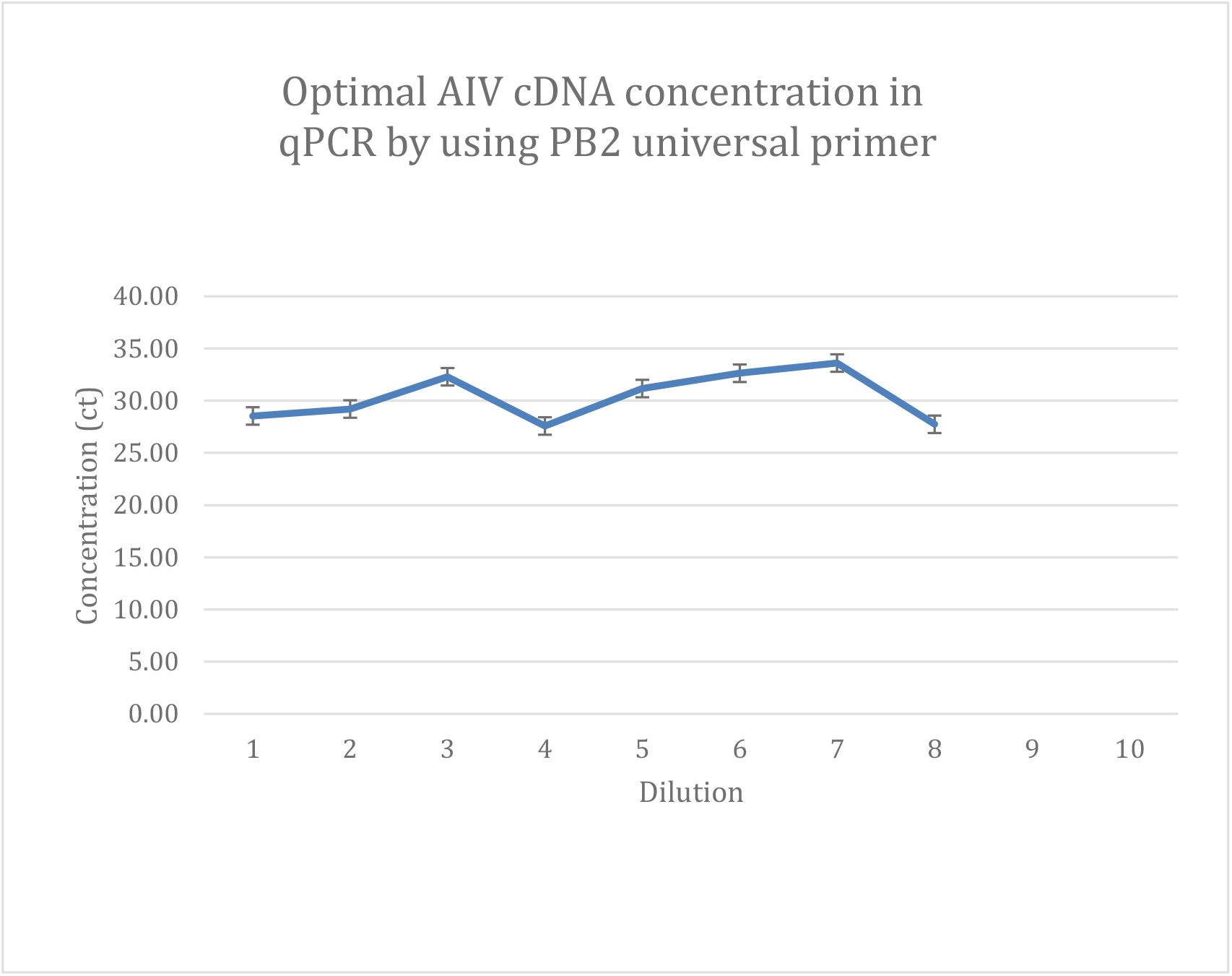
Sensitivity of RRT-PCR with universal primer PB2 for purified AIV sample. The optimal cDNA concentrations of diluted AIV isolates measured with primer set PB2 to measure sensitivity.

**Figure 3.**
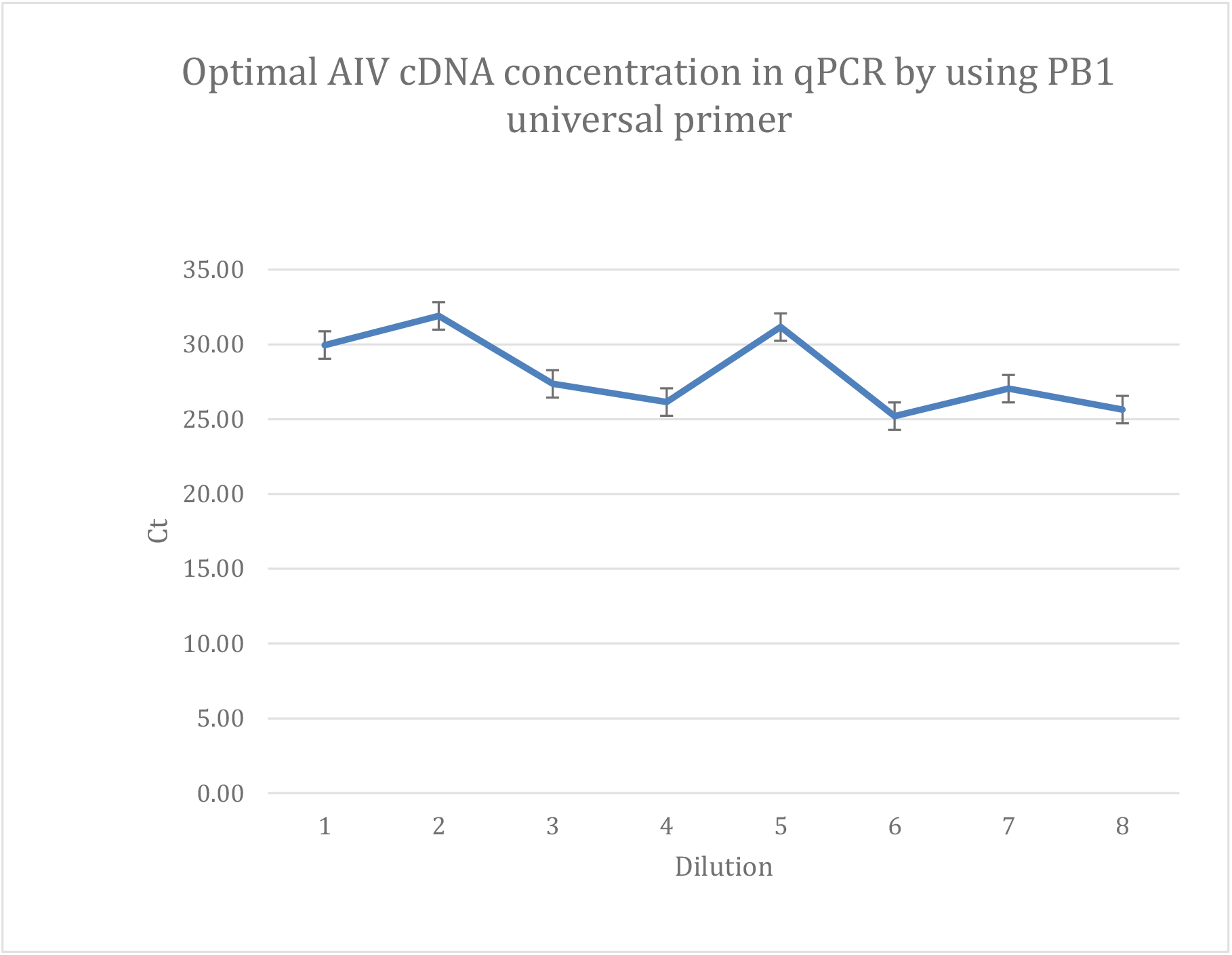
Sensitivity of RRT-PCR with universal primer PB1 for purified AIV sample. The optimal cDNA concentrations of diluted AIV isolates measured with primer set PB1 to measure sensitivity.

Amplification of cDNA from each field sample using RRT-PCR yielded an observed signal after 20 cycles. Intense signals were observed at 28 cycles which indicated a maximum relative florescence unit (RFU) approximately 750 and an average about 350 RFUs demonstrated in Figure 1. RFU values increased detecting higher quantities of DNA. At 40 cycles maximum reads were as high as 3400 RFUs average to over 600 RFUs. Generally, while some signals could be easily observed at 22 cycles and exponentially raised higher in their relative fluorescence, others had a delayed read beginning at 32 cycles later.

To test for sensitivity of universal primers in qPCR, the cDNAs were diluted. The PB2 universal primer set showed a general unusual sensitivity even with increasing dilutions with an average concentration of 30 ct per each dilution. Even at a dilution of 10^8^ the primer was equally sensitive at a lower concentration shown in Figure 2. Similar to primer PB2, PB1 remained consistent across eight dilutions. Starting with 30 ct at 10^1^ dilution it declined on the average to 26 ct at 10^4^ and remained consistent on average through the remaining of the four serial dilutions. This trend remained constant to the highest dilution shown in Figure 2. This indicates an unbiased primer giving an almost equal concentration of the amplified cDNA from the template synthesized.

The optimal AIV cDNA concentration in qPCR universal primer PA, HA, NP, NA and NS showed some unique discrepancies in its sensitivity with dilutions shown in Figures 4,5,6,7 and 9 respectively. Apart from their unusually low yield of amplifications (cDNA from qPCR) they each displayed unusual variability between dilutions. At a dilution of 10^1^ each of the five universal primers (PA, HA, NP, NA, and NS) produced cDNA concentrations below 27.75 ct. In all dilutions the highest concentration of cDNA results revealed a 27.75 ct from the universal primer NP. The lowest AIV cDNA from the five primers came from the PA primer set at a concentration of 22.6 ct at dilution 10^1^. A closer look at the performance of individual primers across the eight serial dilutions can be summarized as follows:

**Figure 4.**
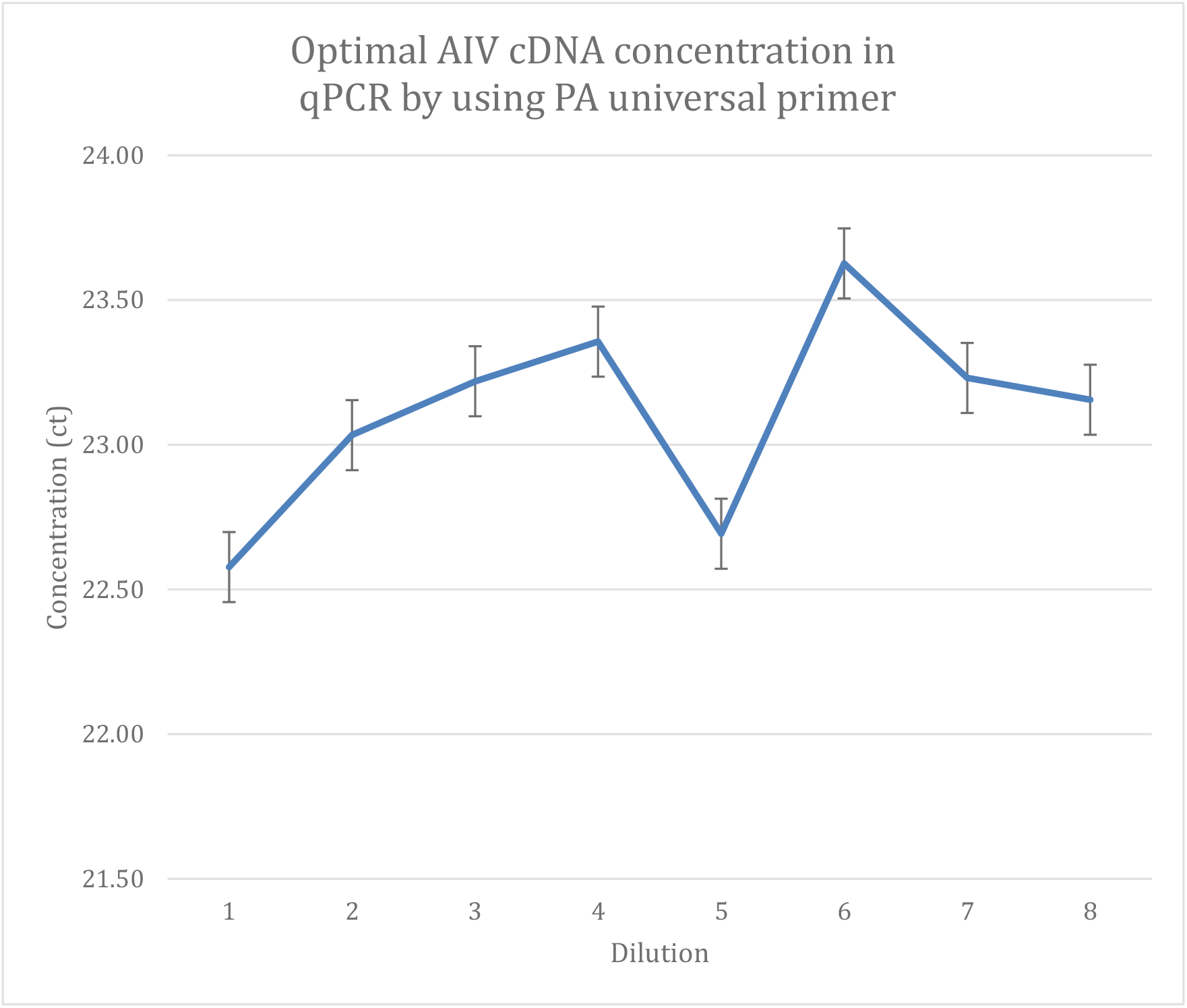
Sensitivity of RRT-PCR with universal primer PA for purified AIV sample. The optimal cDNA concentrations of diluted AIV isolates measured with primer set PA to measure sensitivity.

**Figure 5.**
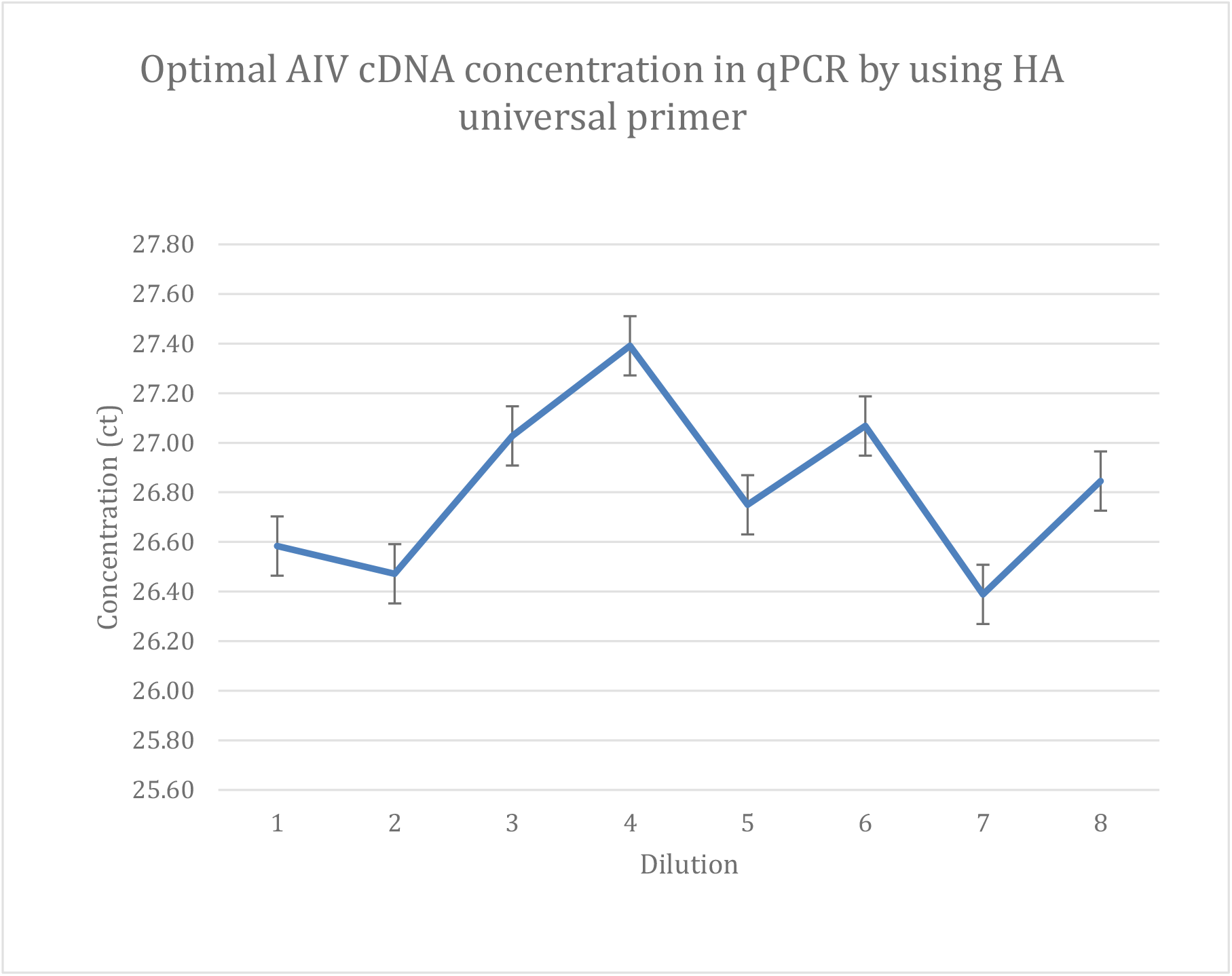
Sensitivity of RRT-PCR with universal primer HA for purified AIV sample. The optimal cDNA concentrations of diluted AIV isolates measured with primer set HA to measure sensitivity.

**Figure 6.**
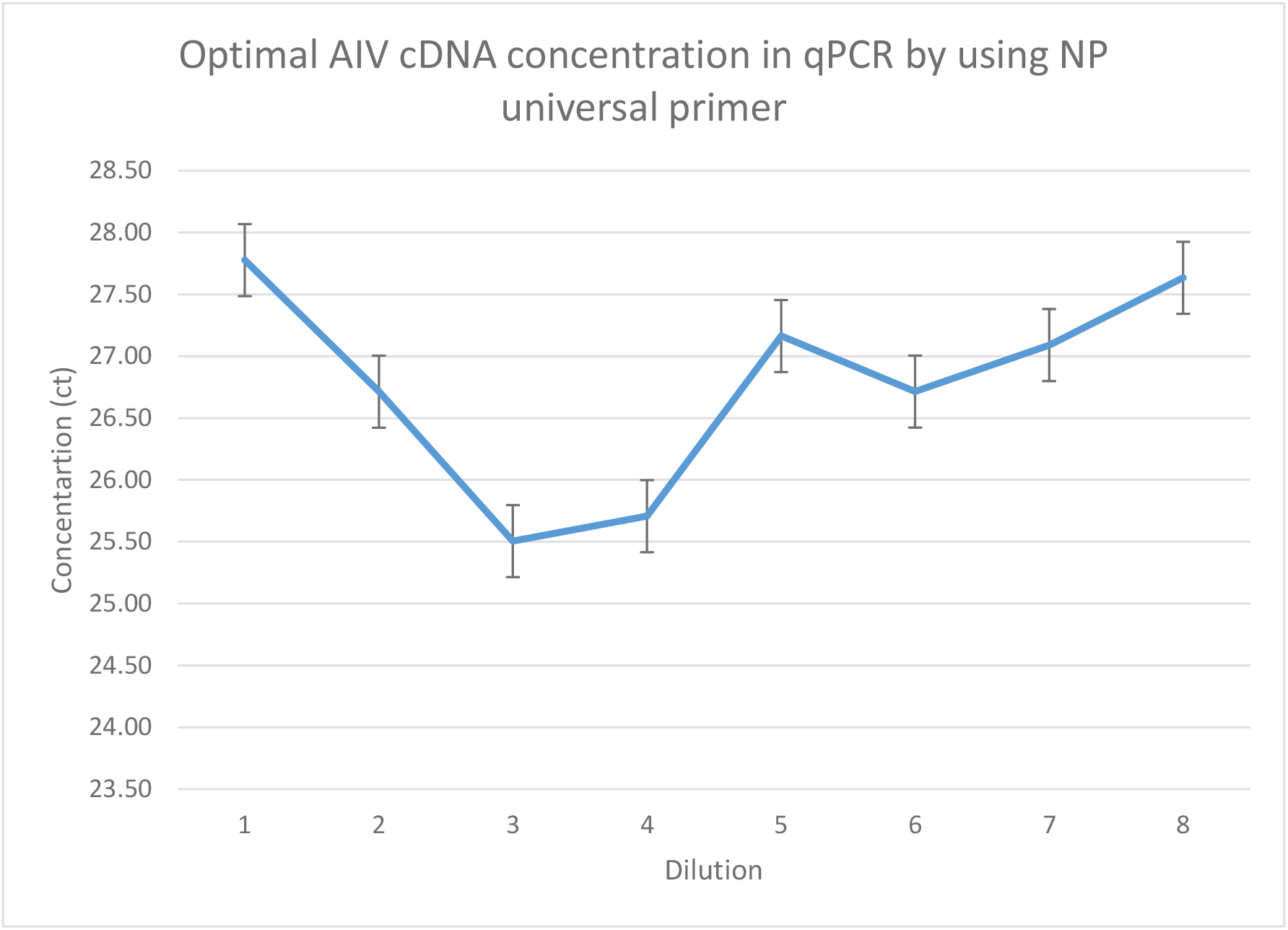
Sensitivity of RRT-PCR with universal primer NP for purified AIV sample. The optimal cDNA concentrations of diluted AIV isolates measured with primer set NP to measure sensitivity.

**Figure 7.**
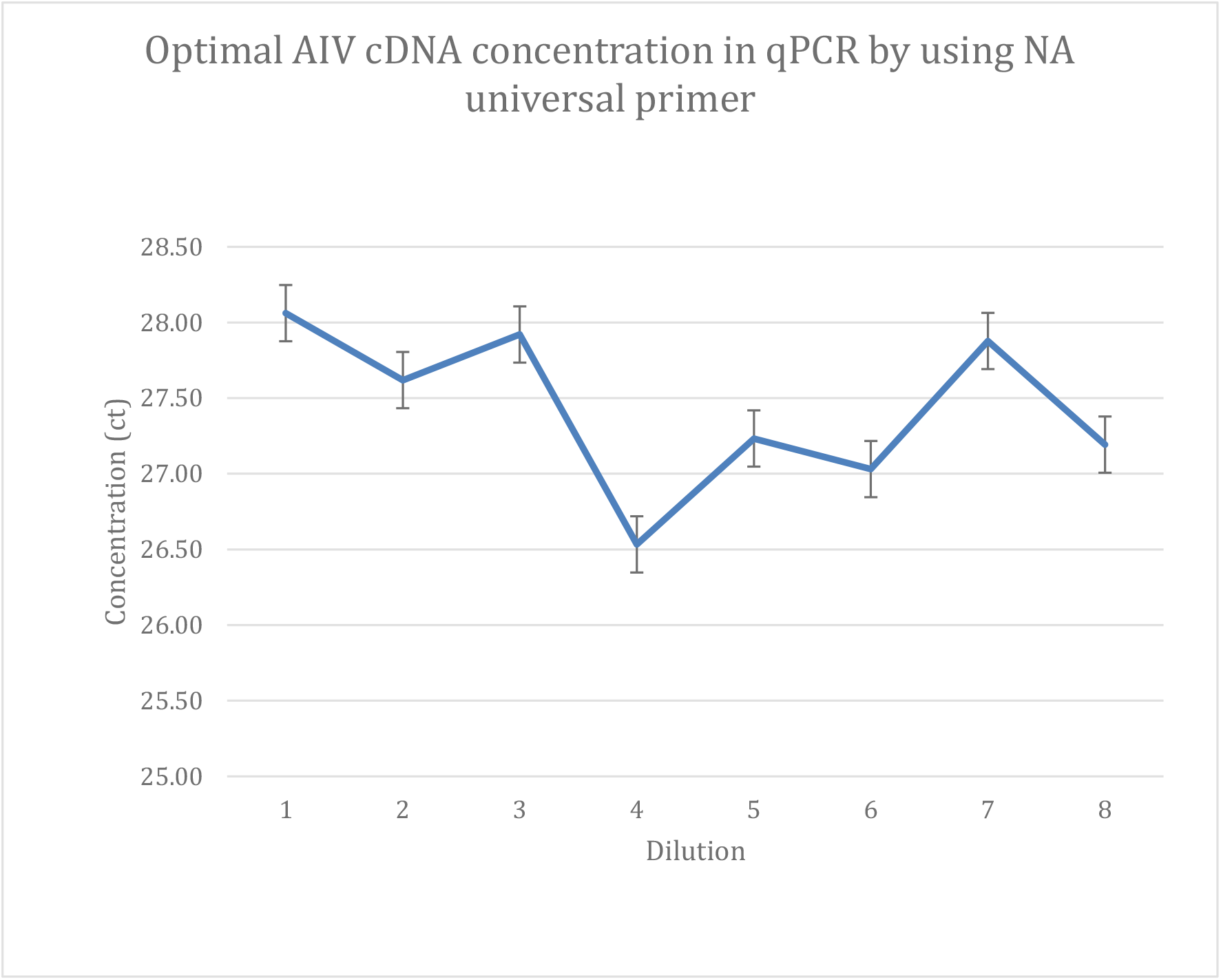
Sensitivity of RRT-PCR with universal primer NA for purified AIV sample. The optimal cDNA concentrations of diluted AIV isolates measured with primer set NA to measure sensitivity.

PA universal primer set in Figure 4 yielded a minimum AIV cDNA at dilution 10^1^ and steadily increased until the product peaked at 23.6 ct in dilution 10^6^ from where it slowly declined to end point at 23.2 ct at 10^8^ dilution.

HA universal primer set showed 26.60 ct and increased with higher dilutions until it peaked at 27.40 ct in dilution 10^4^ from where is slowly declined to stay at 26.80 ct at 10^8^ dilution showing an inverted curve. This is displayed in Figure 5.

NP primer set showed an interesting curve that seemed to fit an ideal scenario showing a higher concentration of 27.75 ct AIV cDNA at lower dilutions and decline with increased dilutions. It remained constant at dilutions 10^3^ and 10^4^ and then increased slowly up to 27.60 ct at 10^8^ dilution displayed in figure 6.

NS primer set showed its highest qPCR product concentration at 10^1^ and steadily declined until the concentration reached its lowest of 24.50 ct. at a dilution of 10^6^. From here its sensitivity increased to result in 25.50 ct and 24.80 ct, respectively at dilutions of 10^7^ and 10^8^.

M universal primer set shown in Figure 8 is produced optimal AIV cDNA concentration averaging 28 ct through all eight serial dilutions. Specifically, at 32 ct at 10^1^ dilution the concentration lowered to 27 ct at 10^2^ and stayed constant at this range in all dilutions. The results of the primer set appeared to be unaffected by dilutions and showed a consistent output of AIV cDNA from all dilutions. This shows that M primer is not affected by concentration gradients. M primer demonstrates great reliability as a primer of choice for identifying and diagnosing Avian Influenza test. As opposed to diluting template concentration in the qPCR discussed in Figure 1. We measured the individual universal primers by serial diluting the primer and then treating them with a constant concentration of templates. Our results revealed that primer dilution did not generally adversely affect the RRT-PCR results.

**Figure 8.**
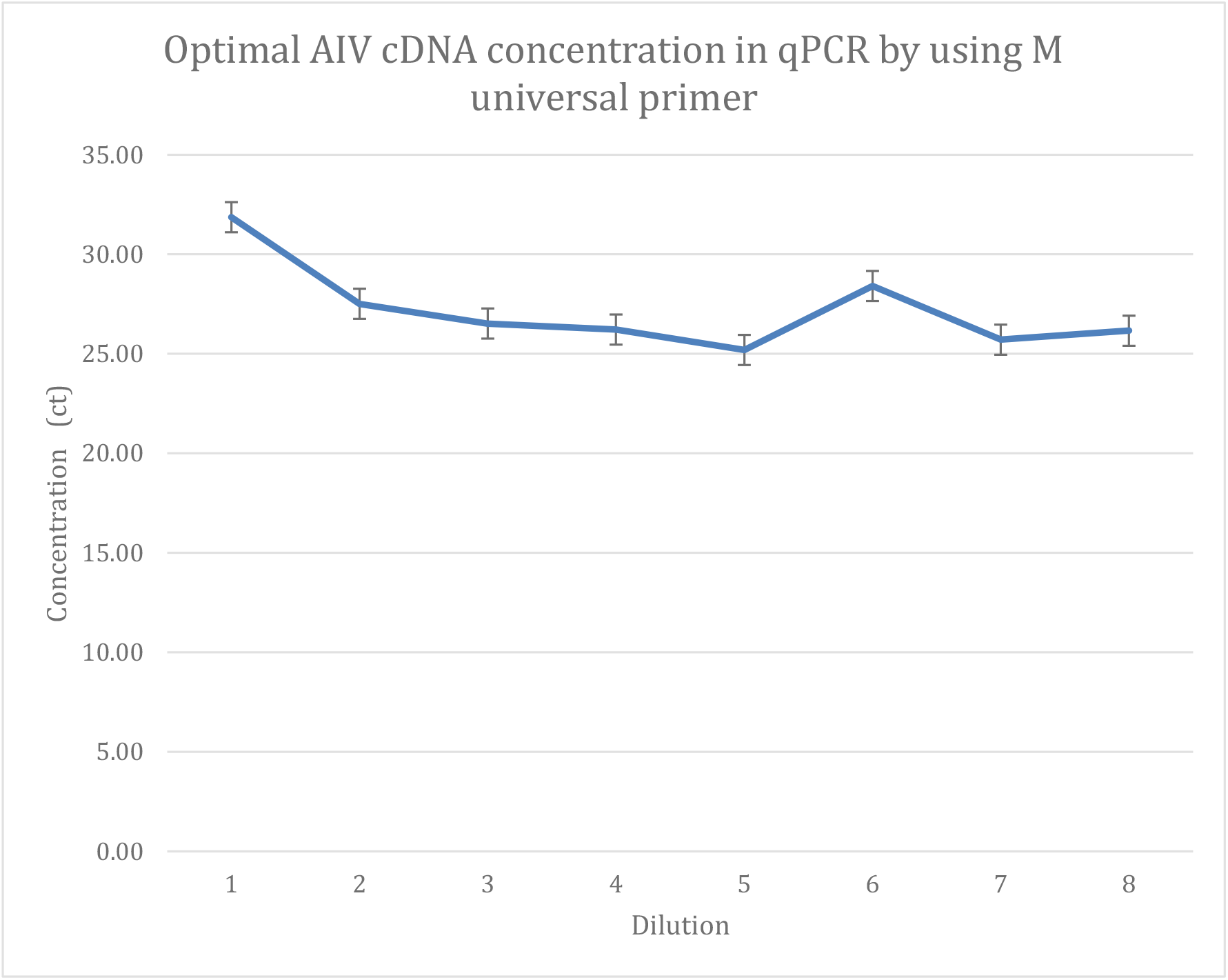
Sensitivity of RRT-PCR with universal primer M for purified AIV sample. The optimal cDNA concentrations of diluted AIV isolates measured with primer set M to measure sensitivity.

### RRT-PCR: The sensitivity of RRT-PCR with serial diluted universal primer

We revealed the template sensitivity of RRT-PCR in table 5. The next step is to further optimize RRT-PCR with serial diluted 8 pairs of universal primer, the results listed in Figure 10-18. Briefly, we selected the optimized cDNA concentration, 8 pairs of universal primer were prepared in ten-fold serial dilutions from 1:10 to 1:10^8^, and real-time PCR was performed as described in the materials and methods. Results of optimal primer summarized in

**Table 5:** Optimized universal primer concentration in RRT-PCR

| Universal Primer | Primer concentration(pmol/μl) |
| --- | --- |
| PB2 | 0.1 |
| PB1 | 0.1 |
| PA | 0.1 |
| HA | 0.01 |
| NP | 1 |
| NA | 1 |
| M | 0.001 |
| NS | 1 |

Optimal primer concentrations measured with positive AIV cDNA isolates. Lowest concentration of 0.001 pmol/ μl and 0.01 detected by primers M and HA.

**Figure 9.**
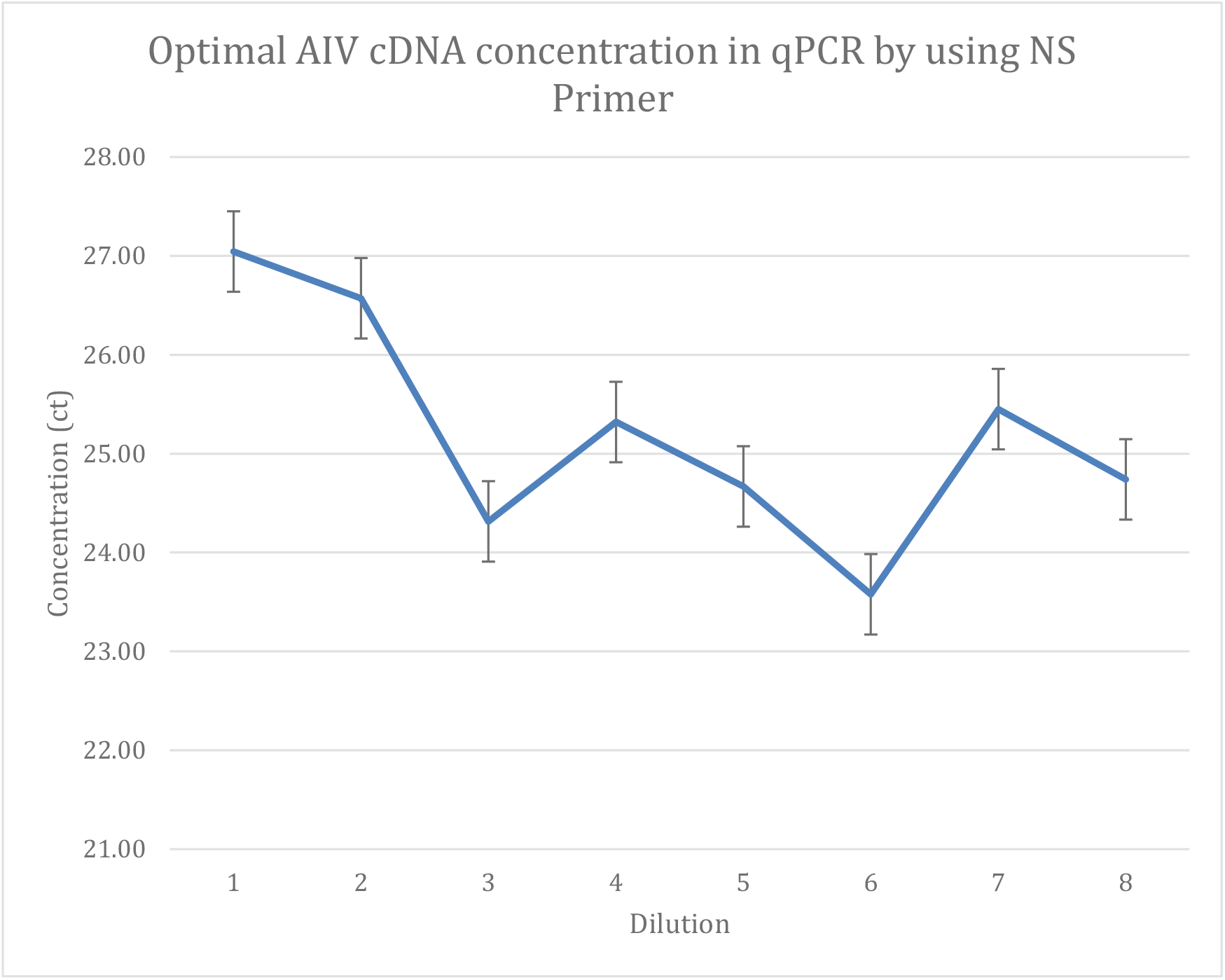
Sensitivity of RRT-PCR with universal primer NS for purified AIV sample. The optimal cDNA concentrations of diluted AIV isolates measured with primer set NS to measure sensitivity.

**Figure 10.**
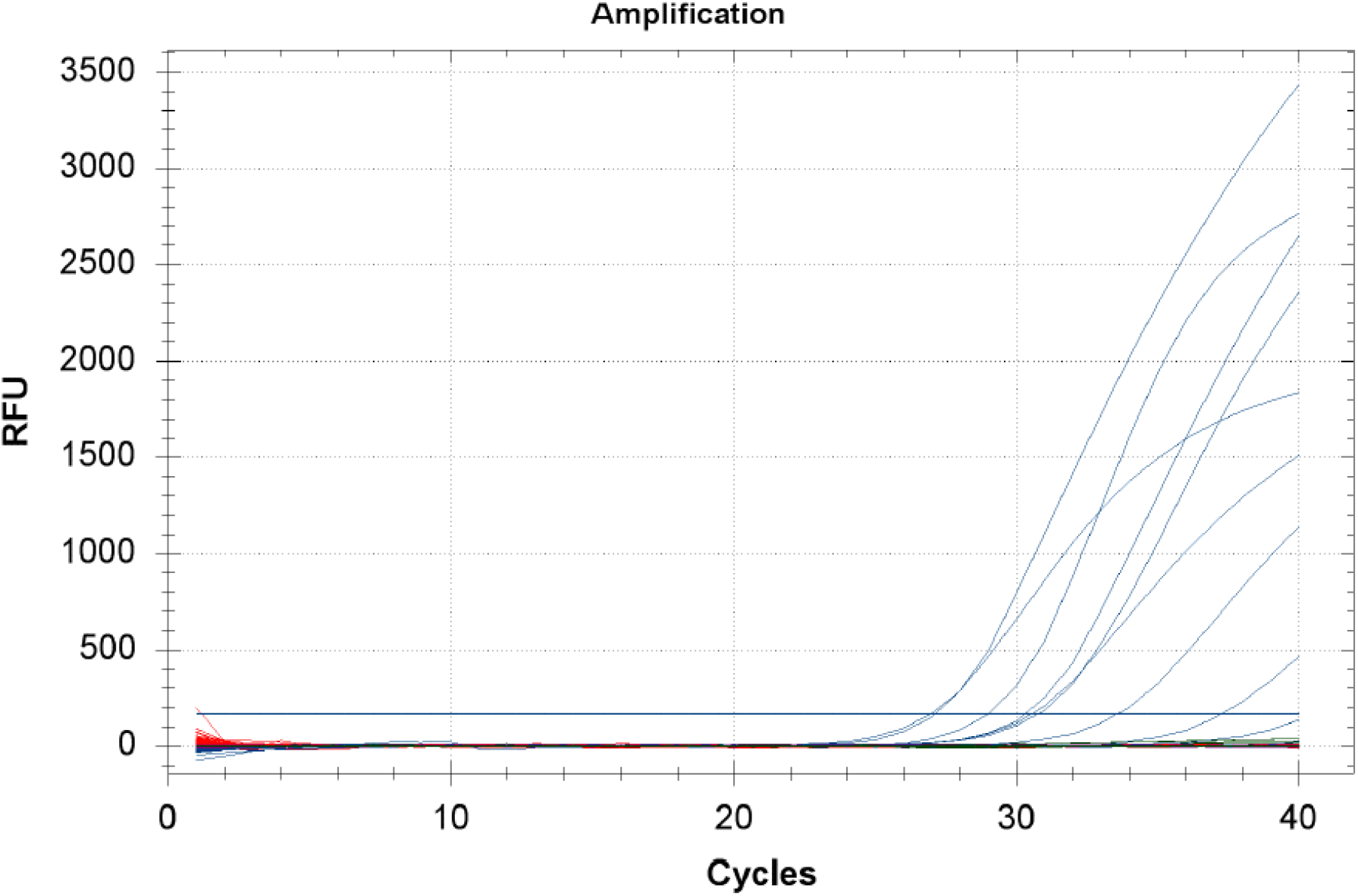
Detection of AIV with RRT-PCR using serial diluted universal primer. Relative Florence Units measured against cycles to determined amplification of diluted universal primers with positive AIV isolates.

In Figure 11, The PB2 gave an output of 34 ct at primer dilution 10^1^ and lowered to 14 ct at 10^3^ primer dilution. At 10^8^ dilution the concentration was at a 24 ct and concentration did not affect the cDNA amplification.

**Figure 11.**
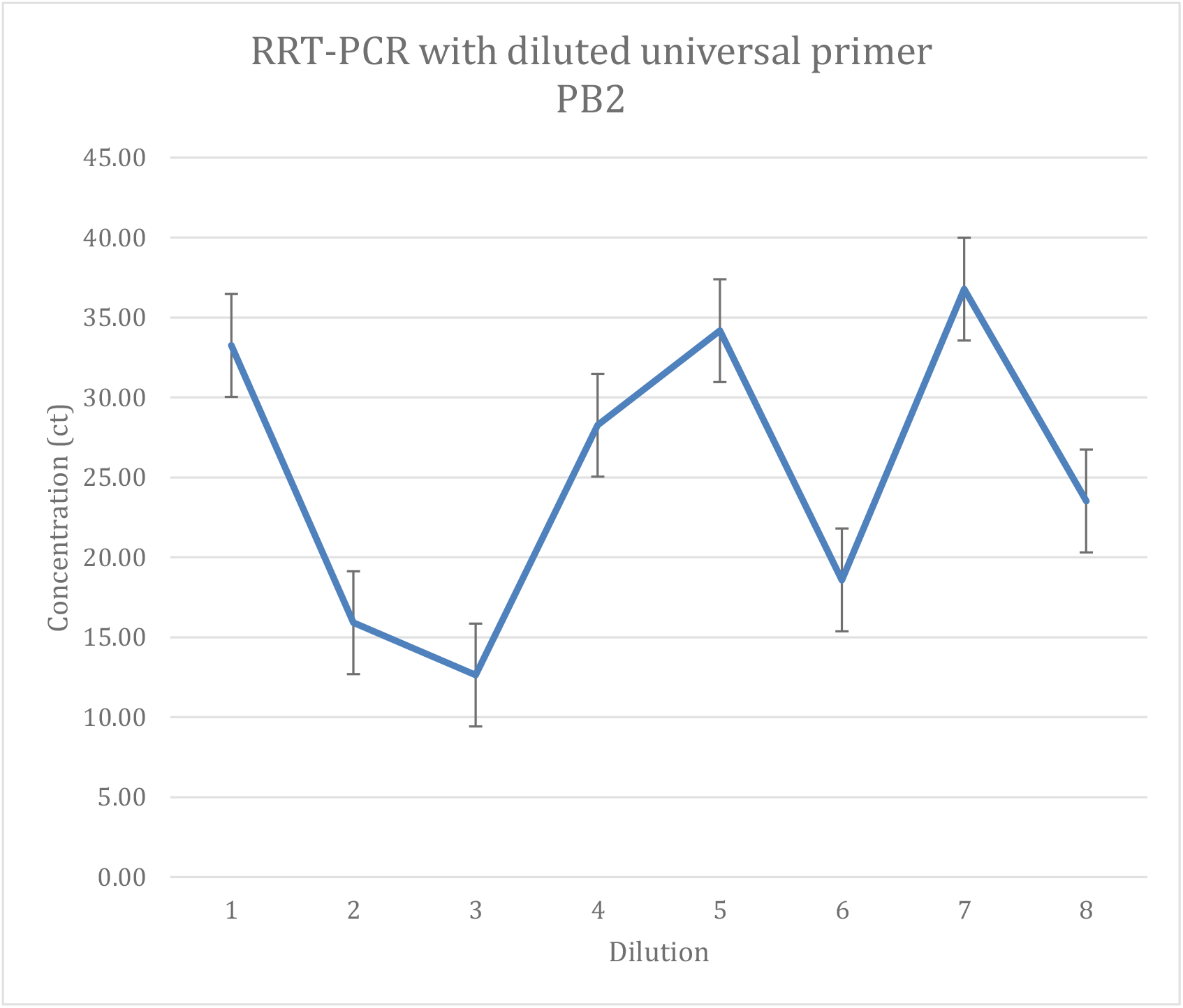
RRT-PCR with diluted universal primer set PB2. Diluted cDNA AIV isolates measured with diluted primer set PB2 to obtain optimal primer concentration to measure sensitivity.

In Figure 12, PB1 primer set showed a rather different trend across concentration gradient. A higher concentration of 10^1^ dilution of primer showed a concentration of 34 ct and steadily declined to a concentration of 13 ct at 10^3^ dilution. The concentration then increased to incrementally produce higher concentrations of 25 ct at a dilution 10^8^.

**Figure 12.**
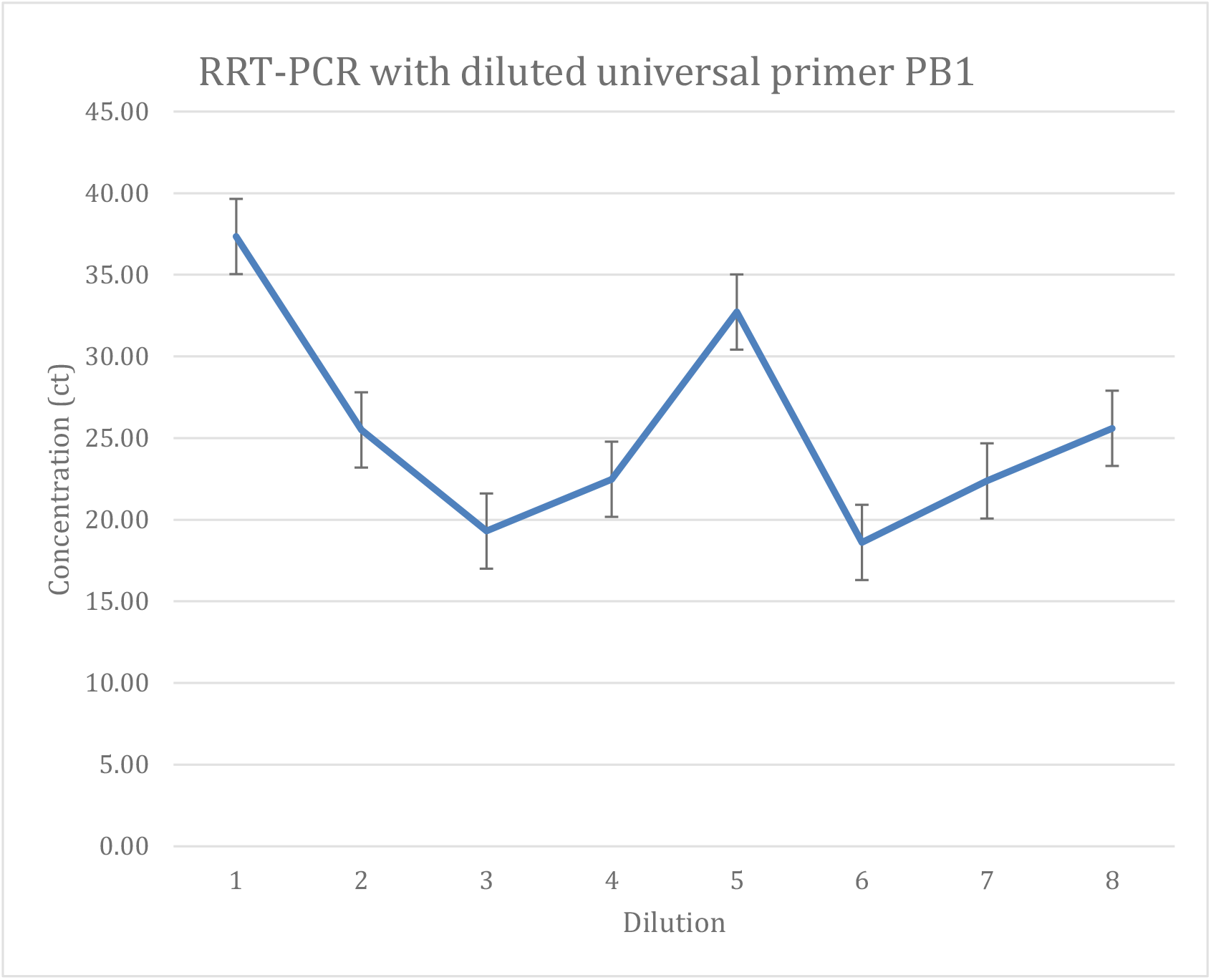
RRT-PCR with diluted universal primer PB1. Diluted cDNA AIV isolates measured with diluted primer set PB1 to obtain optimal primer concentration to measure sensitivity.

PA primer set shown in Figure 13 decreased from 30 ct to 20 ct from 10^1^ to 10^3^ dilutions respectively. It then produced its highest AIV cDNA concentration at 10^4^ then steadily decreased to 5 ct at 10^7^ dilution.

**Figure 13.**
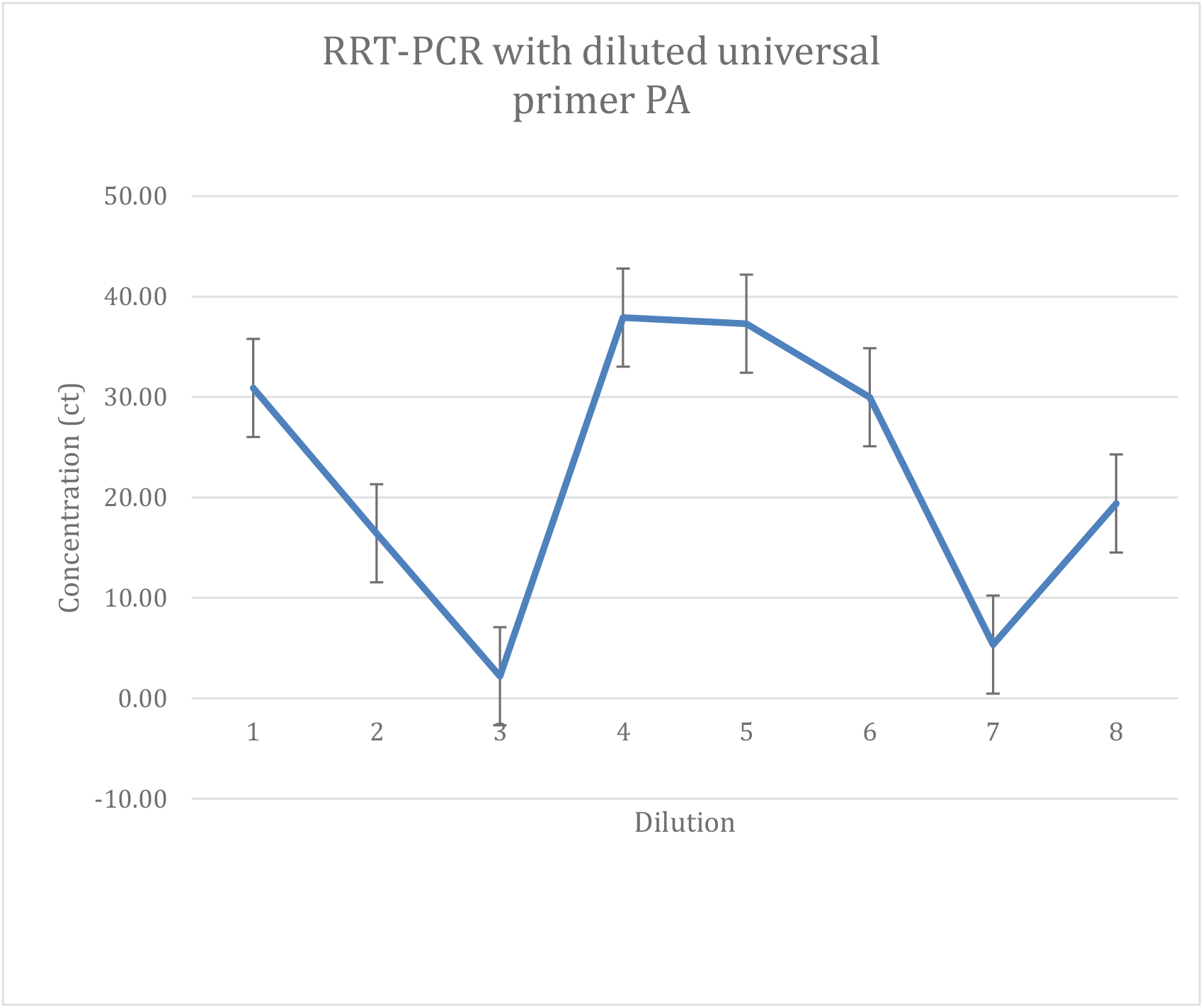
RRT-PCR with diluted universal primer PA. Diluted cDNA AIV isolates measured with diluted primer set PA to obtain optimal primer concentration to measure sensitivity.

In Figure 14, the highest concentration of cDNA was produced at 10^1^ dilution. The lowest concentration was recorded at 2 ct at 10^4^ dilution. The primer at approximate 26 ct of AIV cDNA demonstrated a low concentration of HA primer of 10^5^ to 10^8^ dilutions. In Figure 15, diluting HP in RRT-PCR gave a steady increase with higher dilutions. At 25 ct it continued to the highest concentration of 38 ct at the 10^7^ dilution.

**Figure 14.**
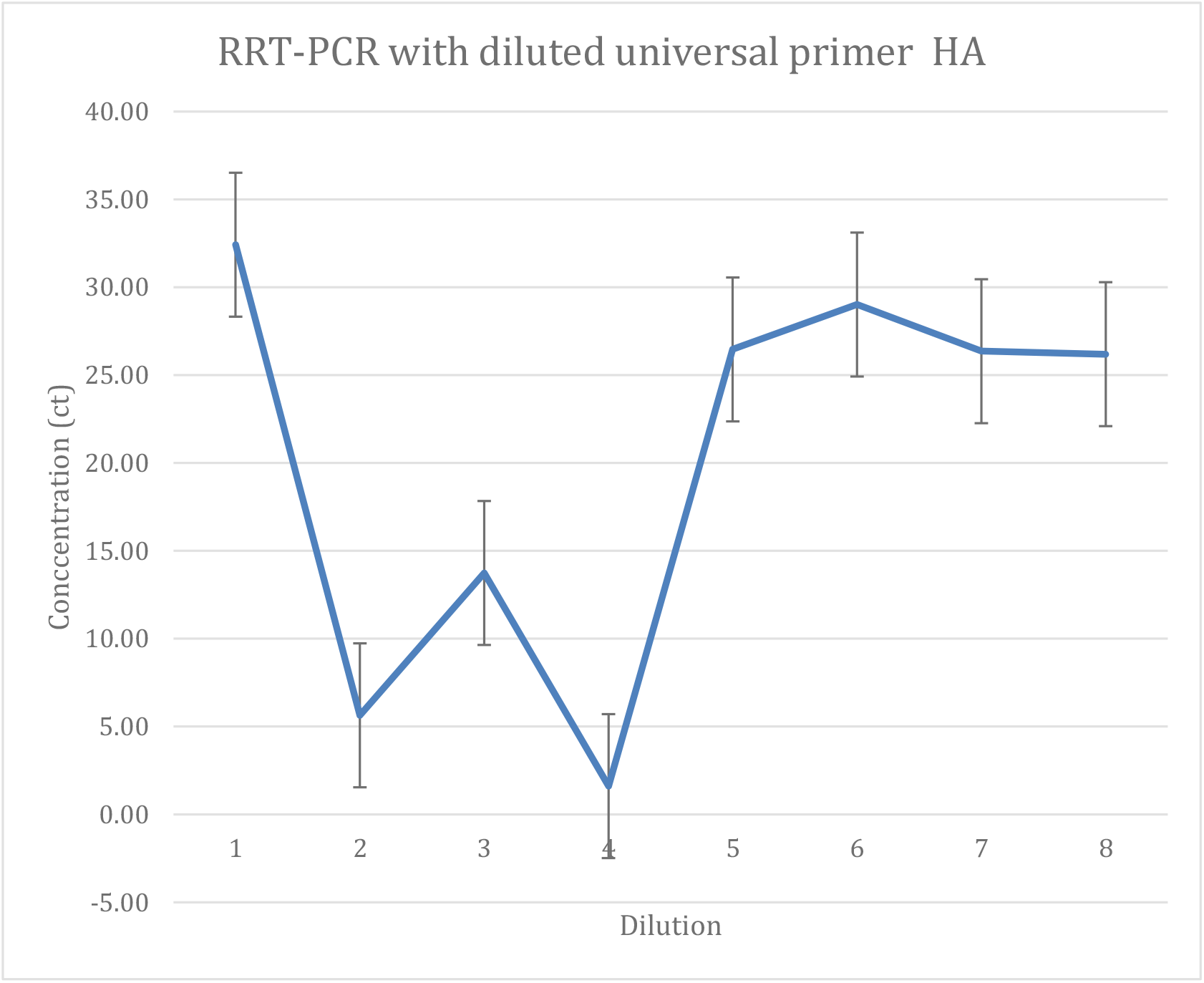
RRT-PCR with diluted universal primer HA. Diluted cDNA AIV isolates measured with diluted primer set HA to obtain optimal primer concentration to measure sensitivity.

**Figure 15.**
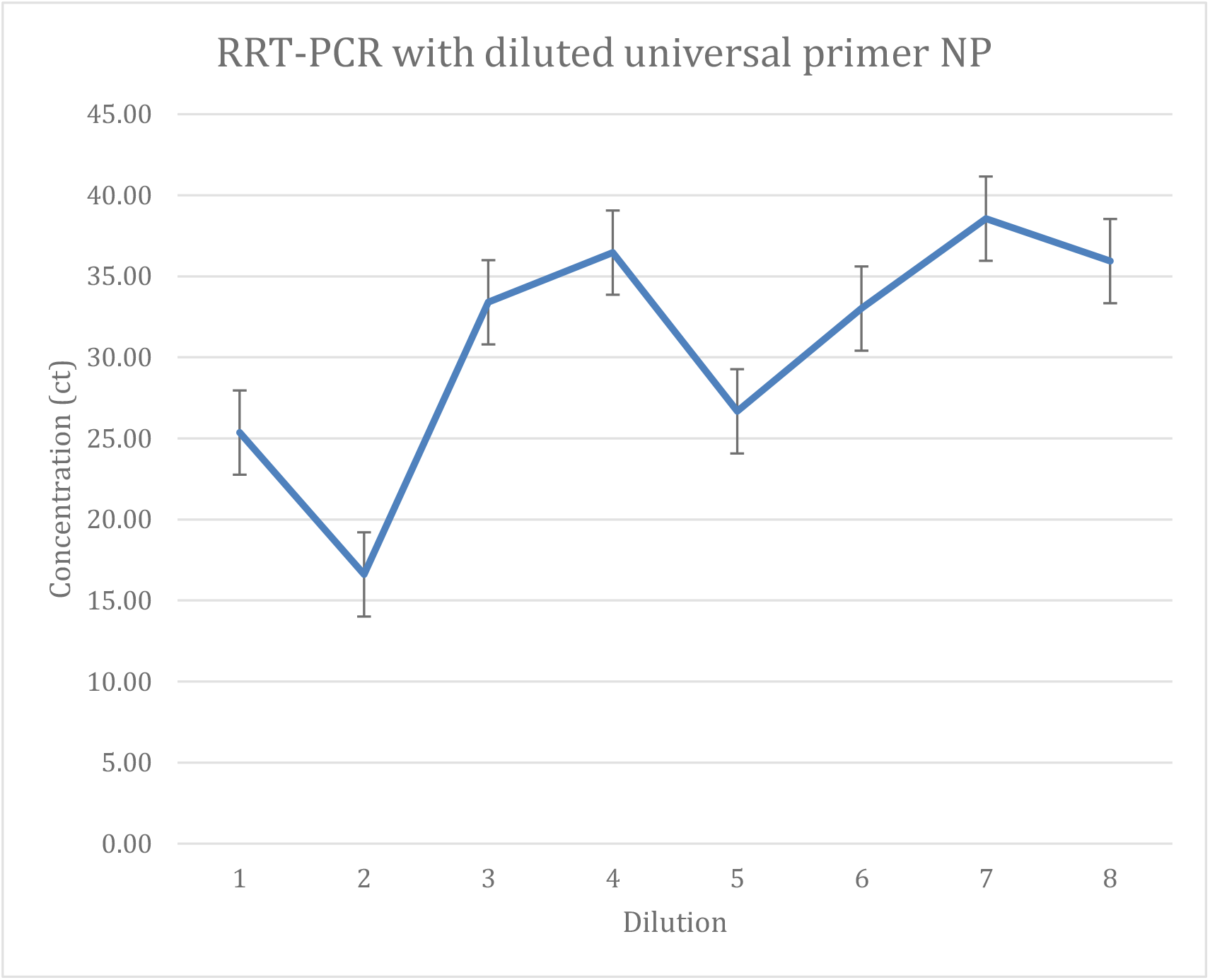
RRT-PCR with diluted universal primer NP. Diluted cDNA AIV isolates measured with diluted primer set NP to obtain optimal primer concentration to measure sensitivity.

NA primer set shown in Figure 16 began at a low of 5 ct at 10^1^ dilution. Then reached a concentration of 35 ct at a 10^3^ dilution. Then trend stayed constant throughout all dilutions until reaching a low of 24 ct at 10^4^. The M universal primer dilution started at a concentration of 33 ct at 10^1^ shown as shown in Figure 17. Generally, primer set M seems to produce a concentration of 25 ct through all dilutions (10^1^ −10^8^). NS however gave a contrasting curve in comparison to the M primer. In figure 18, starting at a dilution of 10^1^ it produced AIV cDNA concentration of 33 ct then reached its lowest at 16 ct at 10^2^ dilution. A concentration of 28 ct at 10^7^ and 10^8^ dilutions of the primers respectively.

**Figure 16.**
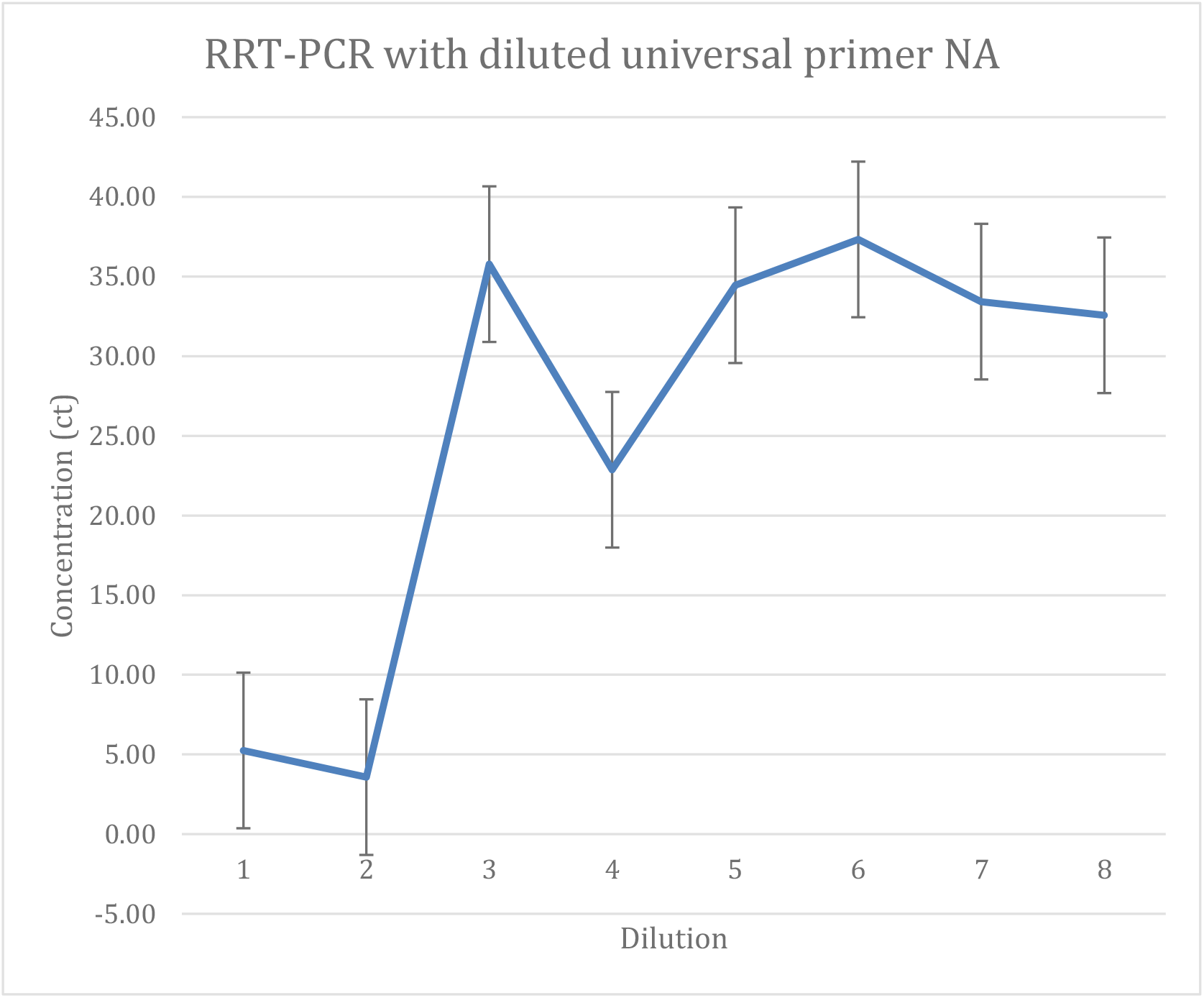
RRT-PCR with diluted universal primer NA. Diluted cDNA AIV isolates measured with diluted primer set NA to obtain optimal primer concentration to measure sensitivity.

**Figure 17.**
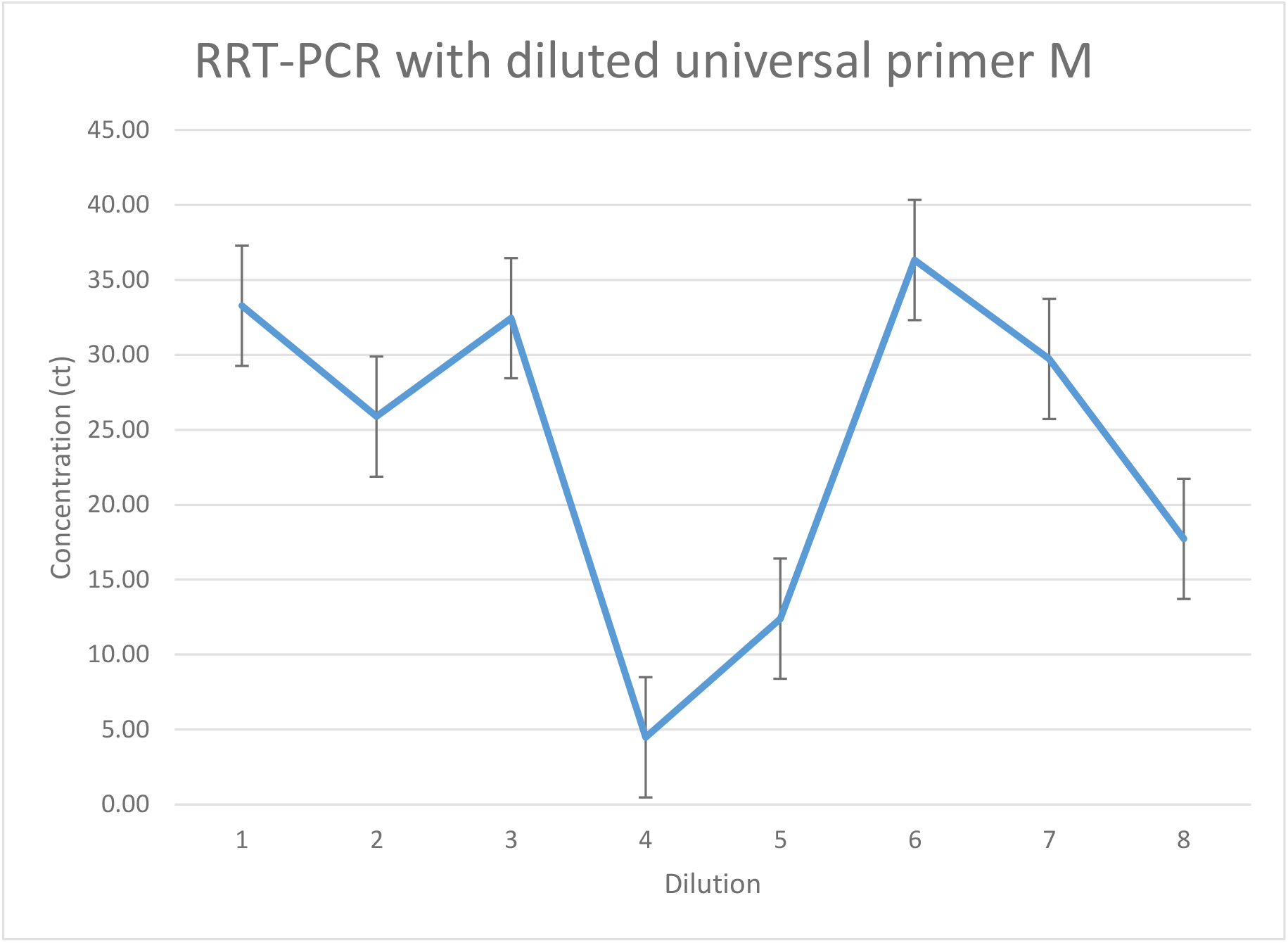
RRT-PCR with diluted universal primer M. Diluted cDNA AIV isolates measured with diluted primer set M to obtain optimal primer concentration to measure sensitivity

**Figure 18.**
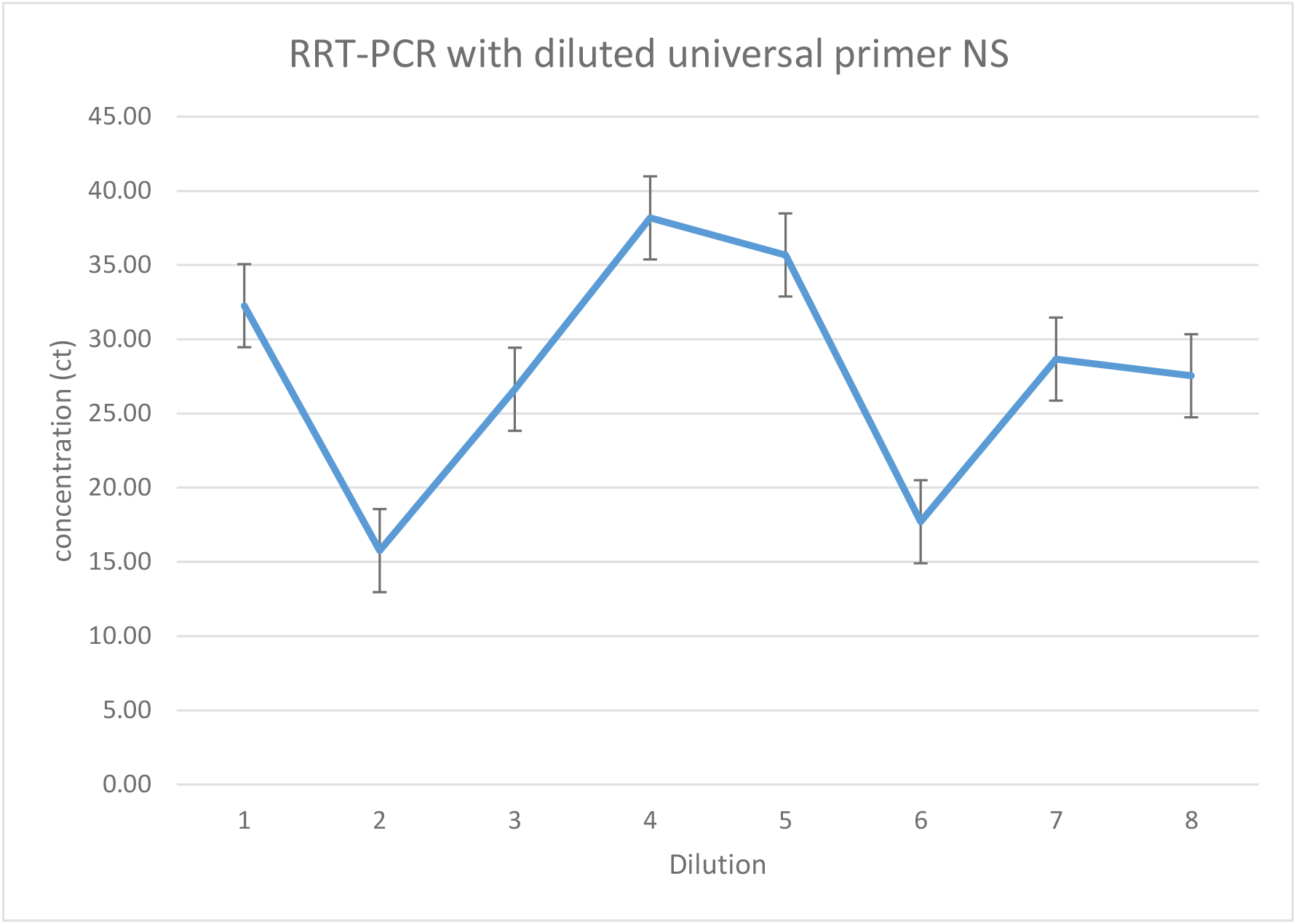
RRT-PCR with diluted universal primer NS Diluted cDNA AIV isolates measured with diluted primer set NS to obtain optimal primer concentration to measure sensitivity.

### RRT-PCR: Detection of AIV in 20 field samples with optimized RRT-PCR

To evaluate the clinical sensitivity of the RRT-PCR under optimized diluted template and universal primer, a total of 20 samples were tested (Figure 19). The results indicated that the RRT-PCR were the best assay for avian fecal samples with universal primer HA in a concentration of 0.01 pmol/μl and sample cDNA concentration 0.0039mg/ml shown in table 6.

**Table 6:** Detection of 20 field samples with optimized RRT-PCR

| Universal Primer | Number of positive samples<br>out of 20 field samples |
| --- | --- |
| PB2 | 4 |
| PB1 | 6 |
| PA | 4 |
| HA | 10 |
| NP | 3 |
| NA | 4 |
| M | 3 |
| NS | 2 |

Primer set HA is identified as the most sensitive primer out of 8 universal primer pairs.

**Figure 19.**
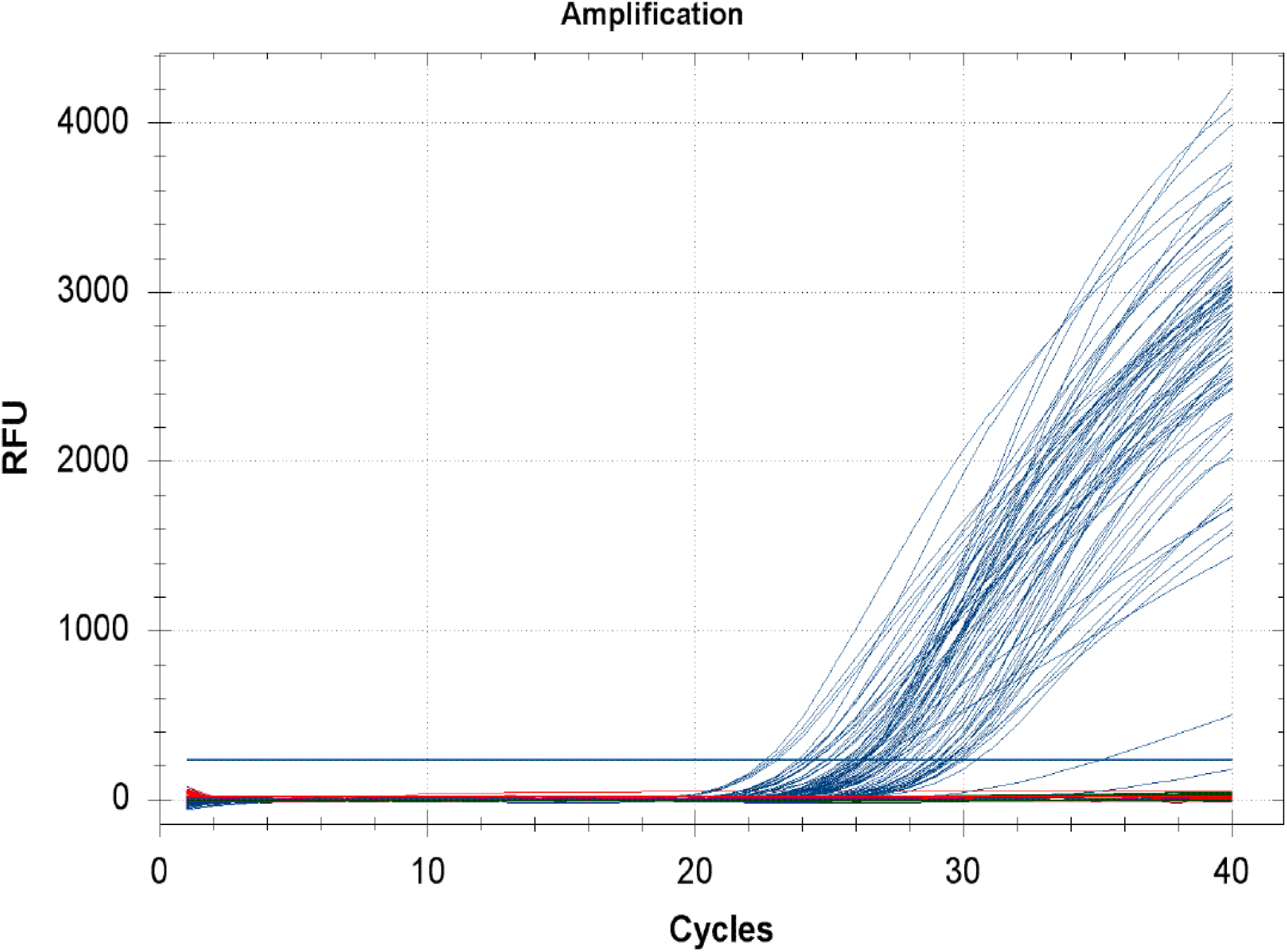
Evaluation of optimized RRT-PCR with field samples. The amplification of 20 field samples measured with 8 universal primer set to identify the most sensitive primer.

## DISCUSSION AND CONCLUSION

Universal primers shown in Table 1 are ideal for this research because they can recognize and bind to any sequence in PCR amplification. AIV viral RNA and cDNA concentrations showed some variations that deviated from expected results shown in Table 2 and Table 3. Normally, RNA concentrations are higher than synthesized cDNA, however there were some discrepancies in which some cDNA field samples were higher. This could have been caused by a number of factors within the spectrophotometer. Cuvettes with residue and dust particles cause light to scatter and can interfere with measuring absorbance accurately of samples. Calibration errors also cause solutions that are outside the required range to cause measuring errors. This typically occurs at lower concentrations (5). The AIV isolates concentrations of cDNA were lower than RNA concentrations. This is expected due to the raw materials of extracted RNA in comparison to cDNA concentrations which is more concentrated with various solutions. In Figures 1, 10 and 19 the number of templates amplified determines Relative Florence Units.

cDNA positive isolates were diluted to measure the primer’s sensitivity at detecting cDNA at very low concentrations. Primer sets PA, HA, NP, and NA were able to detect positive AIV cDNA at low concentrations of 0.0039 mg/ml. These primer sets were most sensitive due to their increased presence of guanine and cytosine in their forward primer. This increases the binding affinity of primers making it easier for primers to recognize, initiate amplification and bind to templates. The number of base pairs in the forward primer is also a useful source when measuring the sensitivity of primers. Hence, the most sensitive primer sets identified in this experiment consists of over 20 to 25 of nucleotides in length. The primers that did not detect cDNA had more adenine and thymine within the forward and reverse primers which resulted in its ability to effectively bind. In Figure 10, there were decreased amplifications of templates this could have been a result of cDNA synthesis activity caused by poor reverse transcriptase. Some other factors include RNA and cDNA degradation during multiple freeze and thaw cycles.

In the identification of the optimal primer concentration, primer set M at 0.001 pmol/µl and HA at 0.01 pmol/µl were able to detect positive cDNA demonstrated in table 5. The binding affinity in both primer sets is increased due to a strong G-C content. In the next experiment, as shown in Figure 19, the collected field samples were collaborated with findings of optimal cDNA and primer concentrations. Primer sets HA and PB1 identified over half of the field samples was positive via RRT-PCR.

In previous research conducted in Dr. Wu’s lab only one primer pair was used in the identification of Avian Influenza. In this study, eight primer pairs with AIV positive isolates determined the optimal concentration of cDNA and primer sets. This was established by using RRT-PCR. Our field samples confirmed laboratory research in identifying positive AIV. Experiments used in this study revealed local wild aquatic birds found at Alabama Shakespeare Park infected with LPAIV and were actively shedding the virus. This is dangerous due to the virus rapid ability to spread. Many visitors including children frequently visit the park and some even feed the birds. As a result, this may increase the risk of infection due to close contact with birds.

Ecological factors, genetic adaptation, and gene assortment have made an emergence of Avian Influenza increasingly possible. These sources, to which can contribute to such a disastrous event, are credited to multiple factors but highly on migratory birds and trade among poultry. In order to prevent an AIV global outbreak it is vital to establish an efficient diagnostic method. Identifying an appropriate assay contributes to infection control and safety measures for human health. Clinical specimens for viral detection are currently limited. Implementing innovative techniques such as using universal primers to identify type A Influenza viruses are imperative. RNA viruses such as AIV has the ability to mutate and crossover to human host cells. Therefore, investigating the evolution and the virus ability to produce new virus strains is critical in preventing, recognizing, and terminating a potential pandemic.

Increased knowledge of the proper testing assay for identifying particular Influenza zoonotic pathogens as well as human transmissible pathogen is vital in diagnosing the virus. This will help scientist to be able to accurately provide a solution in the development of a precise diagnostic assay which is time efficient and economically resourceful.

Surveillance of epidemiology of AIV, in Alabama particularly locally in Montgomery, Al confirmed positive influenza virus in wild aquatic fowl. Fecal samples were collected in winter months when the virus is viable in the environment. The survival period of virus in feces is reportedly 107 days and viable for detection in 15-35°C weather (6). Only fresh samples of stools were collected to increase chances of having virus present. This method was proven to be successful. At the lowest concentration the HA universal primer was revealed as the most sensitive primer out of 20 field samples. The primers successfully detected 10 positive samples shown in table 6. PB1 also demonstrated sensitivity in successfully identifying AIV. The uses of this study are very important due to the increase in the occurrence of Avian Influenza worldwide in poultry and humans. This introduces a major threat for Influenza A pandemic that could pose a threat to both human health and the global economy.

Optimization of sensitive primer used for RRT-PCR will accelerate the diagnosis of AIV and potentially create an accurate therapy method (9,10). Future studies consist of further isolation of Avian Influenza Virus and determining serotype. Further isolation of virus will require a biosafety laboratory level 3 due to risk of inhaling the virus. Collecting field samples from a variety of locations during different times of the year will increase the probability of collecting various strains of Influenza A viruses. Through this research, the RRT-PCR detection assay was optimized and can be used commercially for detection of Influenza A viruses. Optimizing temperatures as well as all primers will also enhance knowledge of primer sensitivity and identification. Optimization of 8 universal primers used in this research is a resourceful time efficient method, that will decrease the rate of false positive results and enable efficient diagnosis.

